# The Hsp40 co-chaperone DNAJC7 regulates polyglutamine aggregation and exhibits context-dependent effects on polyglycine aggregation

**DOI:** 10.1101/2025.08.10.669490

**Authors:** Biswarathan Ramani, Kean Ehsani, Martin Kampmann

## Abstract

Protein-encoding nucleotide repeat expansion diseases, including polyglutamine (polyQ) and polyglycine (polyG) diseases, are characterized by the accumulation of aggregation-prone proteins. In the polyQ diseases, including Huntington’s disease and several spinocerebellar ataxias, substantial prior evidence supports a pathogenic role for mutant polyQ-expanded protein misfolding and aggregation, with molecular chaperones showing promise in suppressing disease phenotypes in cellular and animal models. The goal of this study is to establish a scalable cell-based model to systematically evaluate genetic modifiers of protein aggregation in both polyQ and polyG diseases. We developed FRET-based reporter systems that model polyQ and polyG aggregation in human cells and used them to perform high-throughput CRISPR interference screens targeting all known molecular chaperones. In the polyQ model, the screen identified multiple Hsp70 chaperones and Hsp40 co-chaperones previously implicated in polyQ aggregation and additionally revealed the Hsp40 co-chaperone DNAJC7 as a potent and previously unrecognized suppressor of polyQ aggregation. In contrast, in a FRET-based polyG aggregation model of neuronal intranuclear inclusion disease, CRISPRi screening showed minimal overlap of chaperone modifiers of the polyQ screen. Direct knockdown of DNAJC7 also did not affect polyG aggregation, yet overexpressed DNAJC7 co-localized with both polyQ and polyG aggregates in cells and reduced their aggregation. In addition to establishing new inducible, scalable cellular models for polyQ and polyG aggregation, this work expands the role of DNAJC7 in regulating folding of disease-associated proteins.

## Introduction

An increasing number of neurodegenerative diseases are now recognized to be caused by nucleotide repeat expansions that are in protein-coding regions. Among these are the polyglutamine (polyQ) disorders caused by CAG repeat expansions, including Huntington’s disease (HD), spinocerebellar ataxias (SCA1, 2, 3, 6, 7, 17, and 51), spinobulbar muscular atrophy, and dentatorubropallidoluysian atrophy [1,2]. Similarly, diseases such as neuronal intranuclear inclusion disease (NIID) and fragile X-associated tremor/ataxia syndrome have more recently been linked to GGC repeat expansions that encode polyglycine (polyG) tracts. A common neuropathologic hallmark of these disorders is the accumulation of misfolded, aggregated proteins in the nucleus of neurons, often forming discrete intranuclear inclusions that immunostain with the autophagy adapter protein p62 [3–5].

Strong evidence implicates misfolded protein aggregates as pathogenic drivers in neurodegeneration. In both model systems and human patients, longer CAG repeats correlate with increased protein aggregation and more severe clinical phenotypes. Moreover, molecular chaperones, proteins that assist in folding and refolding of misfolded proteins, have been shown to suppress polyQ aggregation and mitigate toxicity in disease models [6–10]. These findings underscore the importance of the molecular chaperone network (i.e. the chaperome) and the broader proteostasis network in disease pathogenesis, highlighting their potential as therapeutic targets [11,12]. Importantly, molecular chaperones and co-chaperones represent a large and very diverse group of over 300 proteins, with only a subset of chaperones examined as possible modifiers of polyQ aggregation.

Here, we report the development of a Förster Resonance Energy Transfer (FRET)-based human cellular model of polyQ disease that exhibits spontaneous nuclear aggregation along with recapitulating other key features resembling human neuropathology. Moreover, the FRET readout enables high-throughput assessment of aggregation status in individual cells via flow cytometry. Using this model, we show the results of an unbiased CRISPR interference (CRISPRi) screen targeting all known molecular chaperones and co-chaperones, revealing DNAJC7 as a strong modifier of polyQ aggregation. We also developed a comparable FRET-based model for polyG nuclear aggregation, which exhibits similar microscopic features but is notably unaffected by DNAJC7 perturbation. Along with expanding the role of DNAJC7, this study demonstrates the utility of FRET-based CRISPRi screening for uncovering regulators of protein aggregation in repeat expansion disorders.

## Results

### A FRET-based reporter of polyQ protein aggregation in the nucleus

A common neuropathological hallmark of polyQ diseases is the accumulation of mutant proteins in neuronal nuclei, where they form detergent-resistant, p62-positive inclusions. To model polyQ aggregation, we used the C-terminal segment of ataxin-3, which harbors the expanded polyQ tract that causes SCA3 and has been shown in prior studies to be highly aggregation prone [13–17]. To create a model amenable to high-throughput analysis, we developed a FRET-based reporter system, which has previously proven useful for detecting aggregation of different disease-associated proteins by flow cytometry [18–21]. We selected the highly efficient FRET pair mNeonGreen and mScarlet [22].

We used a lentiviral backbone containing a Tet-On 3G promoter to generate constructs for doxycycline-inducible expression of nuclear-localized mNeonGreen or mScarlet, each N-terminally fused to a C-terminal ataxin-3 fragment 79 glutamines (Q79). To generate a stable monoclonal cell line, the constructs were packaged into lentivirus and co-delivered into HEK293T cells containing CRISPR interference (CRISPRi) machinery (Fig. 1A). At 5 days after transduction, we incubated the cells with doxycycline overnight, flow sorted for cells containing both fluorescent proteins, and plated cells by serial dilution to isolate a clonal cell line (Fig. 1A). We selected a clone that we henceforth refer to as the “NLS-FRET-Q79” reporter line. This line was selected among other clones based on flow cytometry showing a tight cluster of cells expressing both fluorescent proteins and the emergence of a population of cells with higher FRET signal that coincided with the appearance of microscopic nuclear puncta.

**Fig. 1.**
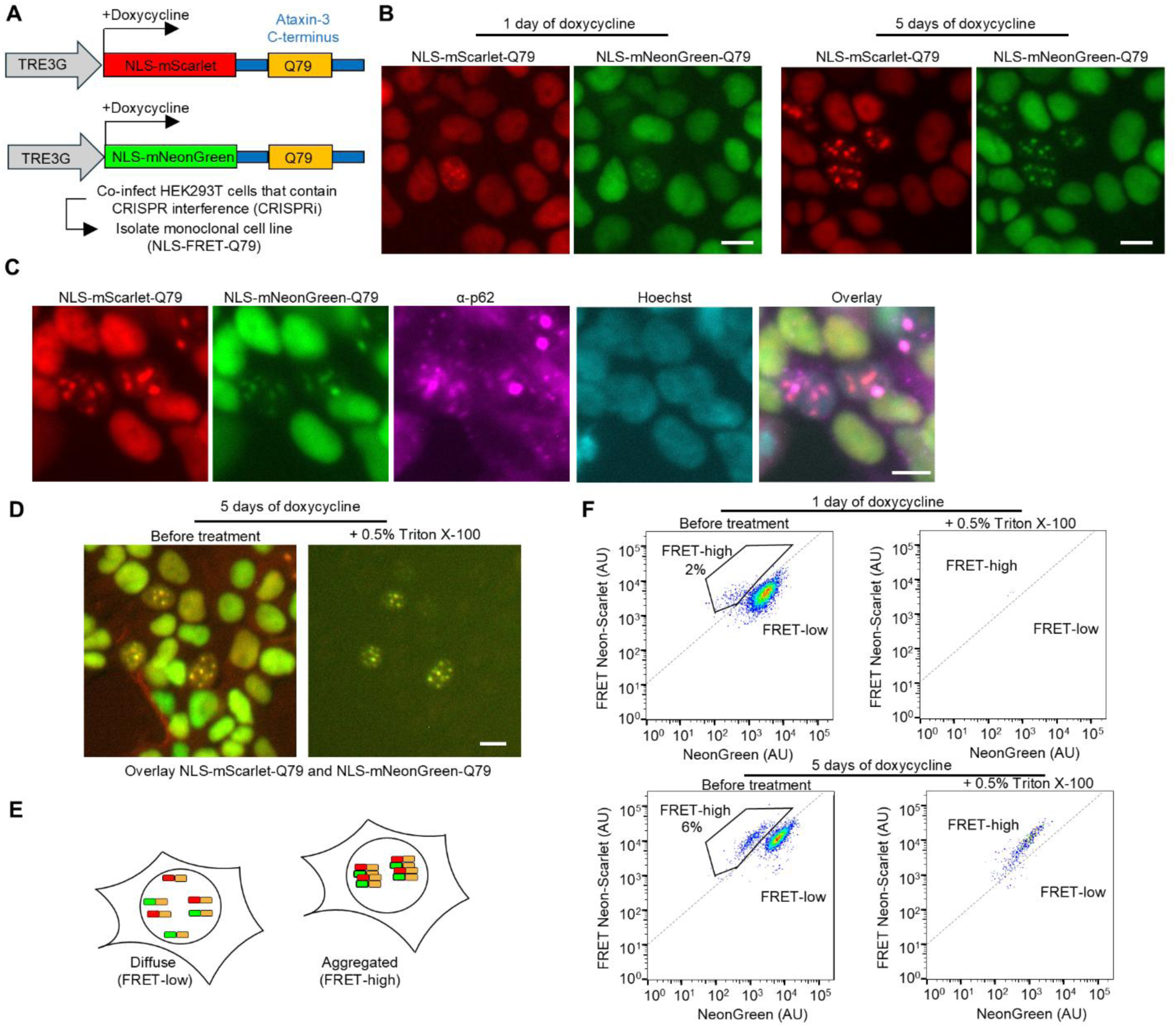
An inducible cell-based model of nuclear, detergent-resistant, p62-positive polyglutamine (polyQ) protein aggregates monitored by a FRET-based reporter. **A**, Schematic of lentiviral constructs for doxycycline-inducible expression of nuclear-localized fluorescent proteins fused to a C-terminal ataxin-3 with 79 glutamines. HEK293T cells engineered with CRISPRi machinery were transduced with these constructs, and clones expressing both fluorescent proteins were selected to generate the NLS-FRET-Q79 cell line. **B**, Fluorescence imaging of NLS-FRET-Q79 cells at one day and five days of doxycycline. **C**, Immunofluorescence staining for endogenous p62 in NLS-FRET-Q79 cells after 5 days of doxycycline. **D**, Fluorescence imaging of NLS-FRET-Q79 cells at 5 days of doxycycline before and after treatment with detergent, in the same field of view. **E**, Schematic illustrating how polyQ aggregation gives rise to a “FRET-high” population observable by flow cytometry. **F**, Flow cytometry plots at one day or five days of doxycycline, before and after treatment with detergent. All scale bars are 10 µm.

In the absence of doxycycline, no expression of fluorescent proteins was detected in the NLS-FRET-Q79 cells. One day after doxycycline induction, most cells exhibited diffuse nuclear fluorescence, while a subset displayed small nuclear puncta (Fig. 1B). By five days of induction, these puncta became more prominent and frequent. The puncta contained both mNeonGreen and mScarlet signals, consistent with polyQ-dependent nucleation and co-aggregation.

Immunofluorescence staining revealed that the nuclear puncta co-localized with endogenous p62/SQSTM1 (Fig. 1C), a hallmark of polyQ inclusions in human disease. Furthermore, treatment of live cells with 0.5% Triton X-100 resulted in the complete loss of diffuse fluorescence, while the nuclear puncta remained intact (Fig. 1D), indicating that these aggregates are detergent-resistant. We did not observe any evident toxicity from inducing expression of the FRET reporter (not shown).

Aggregation of mNeonGreen and mScarlet is predicted to increase FRET efficiency due to their proximity within polyQ aggregates (Fig. 1E). After one day of doxycycline induction, flow cytometry analysis of FRET intensity versus donor mNeonGreen signal shows a dominant single cluster of cells with a small emerging population with increased FRET signal, but nearly all fluorescence was lost upon detergent-treatment (Fig. 1F). In contrast, after five days of doxycycline induction, a more distinct population of cells emerged with even higher FRET signal that persisted even after detergent treatment, consistent with the presence of polyQ aggregates seen by microscopy at this timepoint. We refer to these two populations as “FRET-high” and “FRET-low,” corresponding to aggregated and non-aggregated states, respectively. To confirm that the observed aggregation is polyQ repeat length-dependent, we generated plasmids using a non-pathogenic Q24 length. Transient co-transfection of the FRET pair plasmids containing Q79, but not Q24, led to the formation of nuclear puncta that coincided with an increased FRET signal by flow cytometry, which was also resistant to detergent treatment (Fig. S1). Together, these data demonstrate that the NLS-FRET-Q79 cell line recapitulates key features of nuclear polyQ aggregation observed in human disease and provides a robust platform for high-throughput detection of aggregate-containing cells by flow cytometry.

### A CRISPRi screen identifies DNAJC7 as a suppressor of polyQ aggregation

To identify molecular chaperones that modulate polyQ aggregation, we performed a CRISPRi screen in the NLS-FRET-Q79 cell line using a library of 2,103 sgRNAs targeting 356 genes encoding all known molecular chaperones and co-chaperones (screen workflow shown in Fig. 2A). The screen revealed several candidate modifiers of aggregation, including Hsp70 chaperones, members of Hsp40/DNAJ family of co-chaperones, and several proteasomal subunits (Fig. 2B). The top hit whose knockdown increased FRET signal was *HSPA8*, a constitutively expressed and broadly acting Hsp70 family member involved in ATP-dependent protein folding. In addition, several subunits of the proteasome, including *PSMG4*, *PSMD4*, *PSMC1*, and *PSMD2*, were among the top hits, where knockdown increased FRET signal. To validate this finding, we treated cells with the proteasome inhibitor carfilzomib, which led to a significant increase in the FRET-high population (Fig. 2C).

**Fig. 2.**
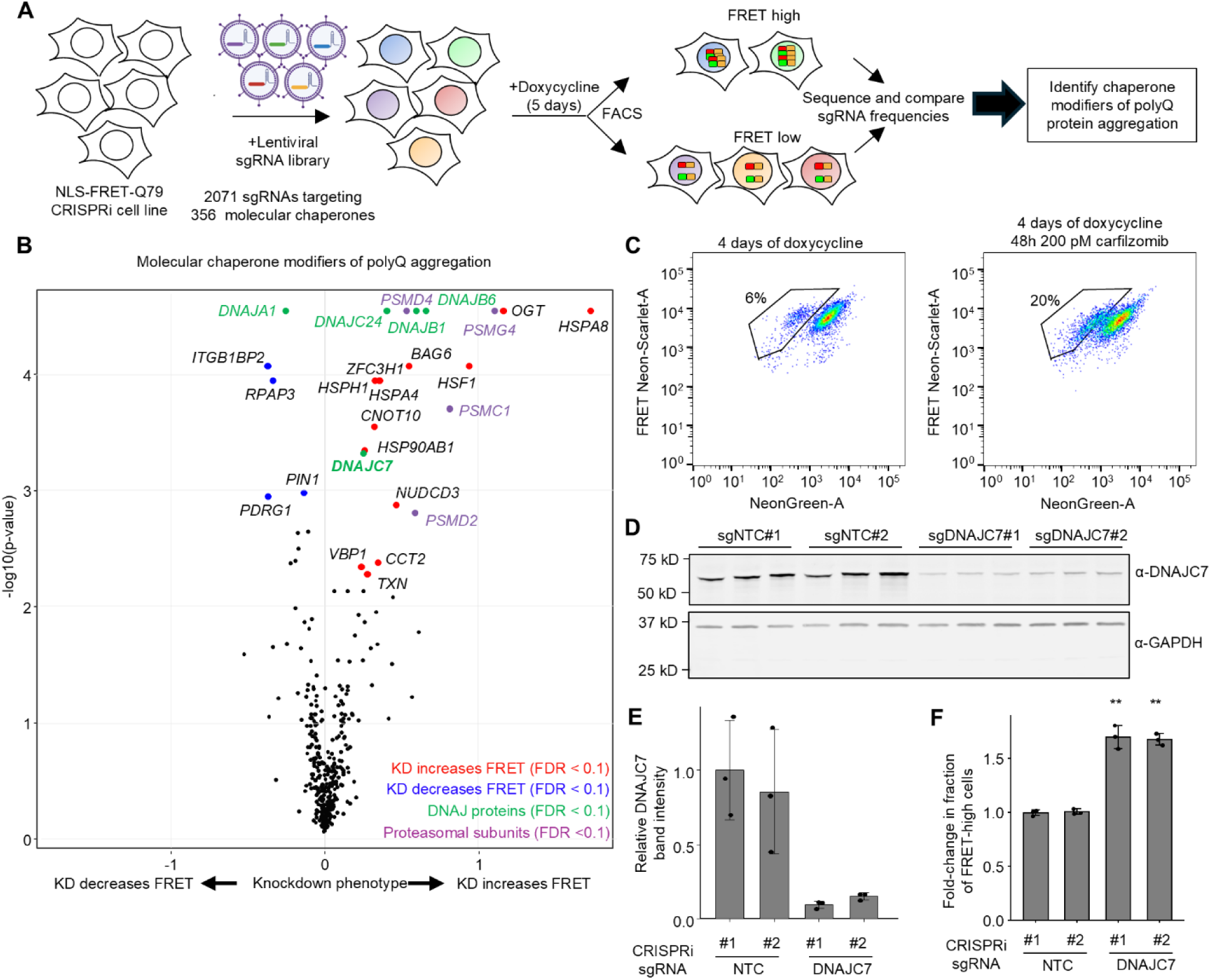
A CRISPRi screen of molecular chaperones identifies DNAJC7 as a modifier of polyQ aggregation. **A**, Workflow of CRISPRi screen to identify molecular chaperone modifiers of polyQ aggregation in NLS-FRET-Q79 cells. Created with BioRender. **B**, Volcano plot CRISPRi screen results, highlighting DNAJ protein and proteasomal subunit hits. **C**, Flow cytometry plots of NLS-FRET-Q79 cells at 4 days of doxycycline with or without proteasomal inhibitor (carfilzomib). **D-E**, Western blot (**D**) and quantification (**E**) of DNAJC7 levels in NLS-FRET-Q79 cells expressing non-targeting (NTC) sgRNAs or sgRNAs targeting DNAJC7. Each lane represents lysate from an independent well; bars and error bars indicate mean ± sd and points represent *n* = 3 wells. Signal intensities were normalized to total protein and then to sgNTC#1. **F**, Results of flow cytometry measuring fold-change in the fraction of FRET-high cells after 5 days of doxycycline in sgRNA^+^ cells transduced with NTC or DNAJC7. Data are presented as fold-change relative to the mean of the sgNTCs for each independent experiment. Bars and error bars represent mean ± sd and points represent *n* = 3 independent experiments. **p < 0.05, one-sample *t*-test compared to sgNTCs (fold-change = 1.0).

Whereas Hsp70s and the proteasome play broad, non-specific roles in protein quality control, the diverse Hsp40/DNAJ family of nearly fifty co-chaperones is thought to confer substrate specificity to Hsp70-mediated refolding [23,24]. Among the top DNAJ family hits in our screen where knockdown increased polyQ aggregation were *DNAJB6*, *DNAJB1*, *DNAJC24*, and *DNAJC7*, while knockdown of *DNAJA1* decreased aggregation. These findings are consistent with prior literature: DNAJB6 and DNAJB1 are well-established suppressors of polyQ aggregation [7,8,25–27], and DNAJA1 knockout cells demonstrate reduced polyQ aggregation [28]. Thus, our screen independently recovers known modifiers of polyQ aggregation, validating the robustness of the approach.

Importantly, in addition to known suppressors, the screen identified *DNAJC7* and *DNAJC24* as previously unexplored hits that significantly increased polyQ aggregation when knocked down. Between these two genes, we prioritized *DNAJC7* for further investigation based on several lines of evidence. Firstly, it is highly expressed in the brain [29]. Secondly, it was recently reported to interact with or modify aggregation of other neurodegenerative disease-associated proteins, including Tau and TDP-43 [30–33]. Thirdly, loss-of-function mutations in *DNAJC7* have been linked to amyotrophic lateral sclerosis [34,35]. Finally, it has been previously identified within polyQ inclusions in mouse neuroblastoma cell models [36,37]. In contrast, while *DNAJC24* was a stronger hit, it has low brain expression and no known links to neurological disease.

To validate the effect of *DNAJC7* knockdown, we generated NLS-FRET-Q79 lines stably expressing either two different non-targeting control (NTC) sgRNAs or sgRNAs targeting *DNAJC7*. Western blotting confirmed efficient knockdown of DNAJC7 protein (Fig. 2D and E).

Importantly, flow cytometry showed a significant increase in the fraction of FRET-high cells upon *DNAJC7* knockdown (Fig. 2F, gating strategy in Fig. S2). These findings confirmed the predictive capacity of the screen and that DNAJC7 is an endogenous suppressor of polyQ aggregation, at least in the context of this FRET reporter system.

### DNAJC7 suppresses mutant HTT exon 1 aggregation and co-localizes with inclusions in cells

To determine whether DNAJC7 modifies aggregation in other polyQ disease contexts, and to exclude the possibility that its effects are specific to the nucleus or the specific FRET pair fluorescent proteins, we generated an independent reporter CRISPRi cell line expressing doxycycline-inducible eGFP-tagged huntingtin exon 1 with 72 glutamines (GFP-HTTex1-Q72) (Fig. 3A). Unlike the NLS-FRET-Q79 construct, HTTex1 naturally contains an N-terminal nuclear export signal, resulting in primarily cytoplasmic aggregation in HEK293T cells [38,39].

**Fig. 3.**
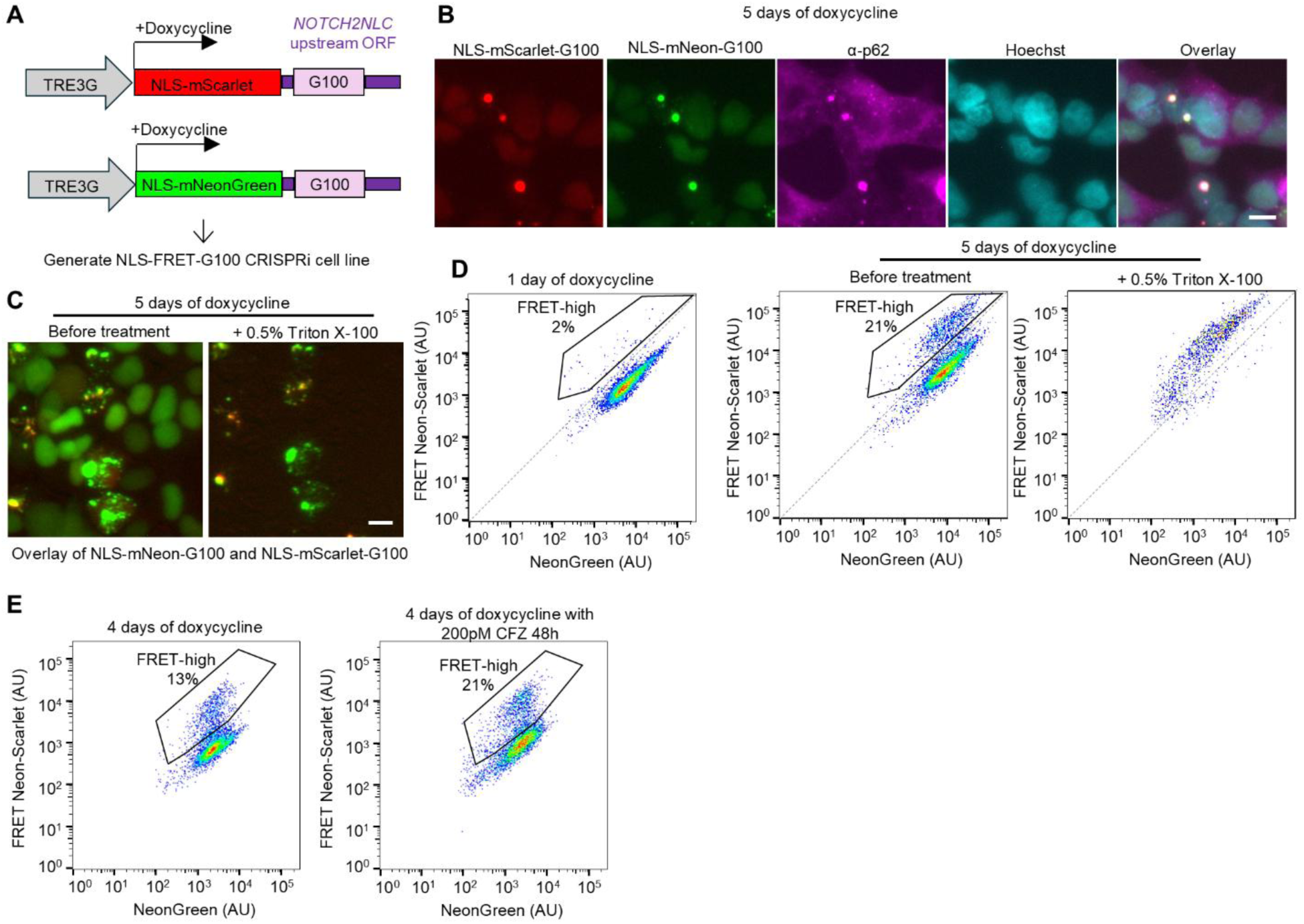
DNAJC7 suppresses mutant HTTex1 aggregation and co-localizes with inclusions. **A**, Schematic of a lentiviral construct encoding a doxycycline-inducible eGFP-tagged fragment of Huntingtin exon 1 containing 72 glutamines (GFP-HTTex1-Q72), used to generate a monoclonal HEK293T CRISPRi cell line. **B**, Fluorescence micrograph of GFP-HTTex1-Q72 cells at 7 days of doxycycline treatment before and after treatment with Triton X-100. **C**, Results of flow cytometry experiments measuring the fraction of total events containing detergent-resistant GFP^+^ cells comparing those stably transduced with NTC or DNAJC7-targeting sgRNAs and normalized to the mean of NTCs. Data represent mean ± sd from *n* = 4 independent experiments. **p < 0.05, one-sample t-test compared to sgNTCs (fold-change = 1.0). **D**, Fluorescence micrographs of HEK293T cells 48 h after transient co-transfection of GFP-HTTex1-Q72 and either mTagBFP2 (BFP) alone or BFP-tagged DNAJC7. Arrows indicate cytoplasmic aggregates. **E**, Flow cytometry plotting GFP-height versus GFP-Width (pulse shape analysis) of HEK293T cells transfected with GFP-HTTex1-Q72 for 48h. The detergent-resistant population of cells are designated as aggregate-positive (Agg^+^). **F**, Results of flow cytometry experiments measuring Agg^+^ cells in HEK293T cells 48h after transient co-transfection with GFP-HTTex1-Q72 and either BFP or BFP-DNAJC7. Bars and error bars represent mean ± sd from *n* = 4 independent biological replicates (indicated by color), with each performed in technical triplicate. *p < 0.05 by Student’s *t*-test using mean values of technical replicates. Scale bars: 10 µm.

After seven days of doxycycline induction, we observed detergent-resistant cytoplasmic GFP+ inclusions in a small subset (∼1-2%) of cells (Fig. 3B). CRISPRi knockdown of *DNAJC7* in the GFP-HTTex1-Q72 cell line significantly increased the fraction of detergent-insoluble GFP+ aggregates, as measured by flow cytometry (Fig. 3C, gating strategy Fig. S2), indicating that DNAJC7 suppresses aggregation in the cytoplasm of a different polyQ protein.

We next sought to determine whether DNAJC7 co-localizes with HTTex1 aggregates. However, attempts to visualize endogenous DNAJC7 by immunostaining were unsuccessful due to the lack of a suitable antibody. To address this, we performed transient co-transfection experiments in HEK293T cells using GFP-HTTex1-Q72 together with either mTagBFP2 (BFP) alone or BFP-tagged DNAJC7. As expected, GFP-HTTex1-Q72 formed cytoplasmic aggregates. Notably, BFP-DNAJC7, but not BFP alone, co-localized with a subset of these aggregates (Fig. 3D), suggesting an interaction between DNAJC7 and aggregated HTTex1.

To assess whether DNAJC7 overexpression could reduce HTTex1 aggregation, we used flow cytometry pulse-shape analysis, which distinguishes aggregates based on a characteristic high-intensity and narrow-width fluorescence signal [27,40]. At 48 h post-transfection, coinciding with the appearance of aggregates by microscopy, we observed a distinct population of cells with high and narrow GFP signal, designated as Agg⁺ (aggregate-positive), which persisted following detergent treatment (Fig. 3E). Compared to BFP control, overexpression of BFP-DNAJC7 significantly reduced the fraction of Agg⁺ cells (Fig. 3F), indicating that DNAJC7 suppresses aggregation of mutant HTTex1 when overexpressed. BFP-DNAJC7 did not reduce the total number of GFP-positive cells compared to BFP (Fig. S3), indicating similar transfection efficiencies.

### A FRET-based reporter for polyG aggregation reveals detergent-resistant, p62-positive nuclear inclusions

Given the emerging role of DNAJC7 as a modifier of multiple disease-associated protein aggregates, we next sought to examine its relevance in a distinct repeat expansion disorder. We focused on neuronal intranuclear inclusion disease (NIID), which is caused by an expanded polyG tract encoded within the upstream open reading frame of the *NOTCH2NLC* (uN2C) gene and, like the polyQ diseases, is pathologically characterized by the presence of p62-positive nuclear inclusions [41,42].

Using a parallel approach to our polyQ model, we developed a clonal HEK293T CRISPRi cell line containing a doxycycline-inducible, nuclear-localized FRET reporter expressing uN2C containing 100 glycines (designated NLS-FRET-G100) (Fig. 4A). Five days after doxycycline induction, a subset of cells exhibited nuclear puncta positive for both fluorophores, which co-localized with p62 (Fig. 4B). These puncta were resistant to Triton X-100 treatment, indicating that they form detergent-insoluble aggregates (Fig. 4C). Flow cytometry revealed a progressive increase in the FRET-high population over time, which was maintained after detergent treatment (Fig. 4D). Inhibition of the proteasome with carfilzomib further increased the FRET-high population (Fig. 4E). By transient transfection experiments, we confirmed that uN2C-G100, but not uN2C-G12 expression led to the formation to detergent-resistant FRET-high cells (Fig. S4)

**Fig. 4.**
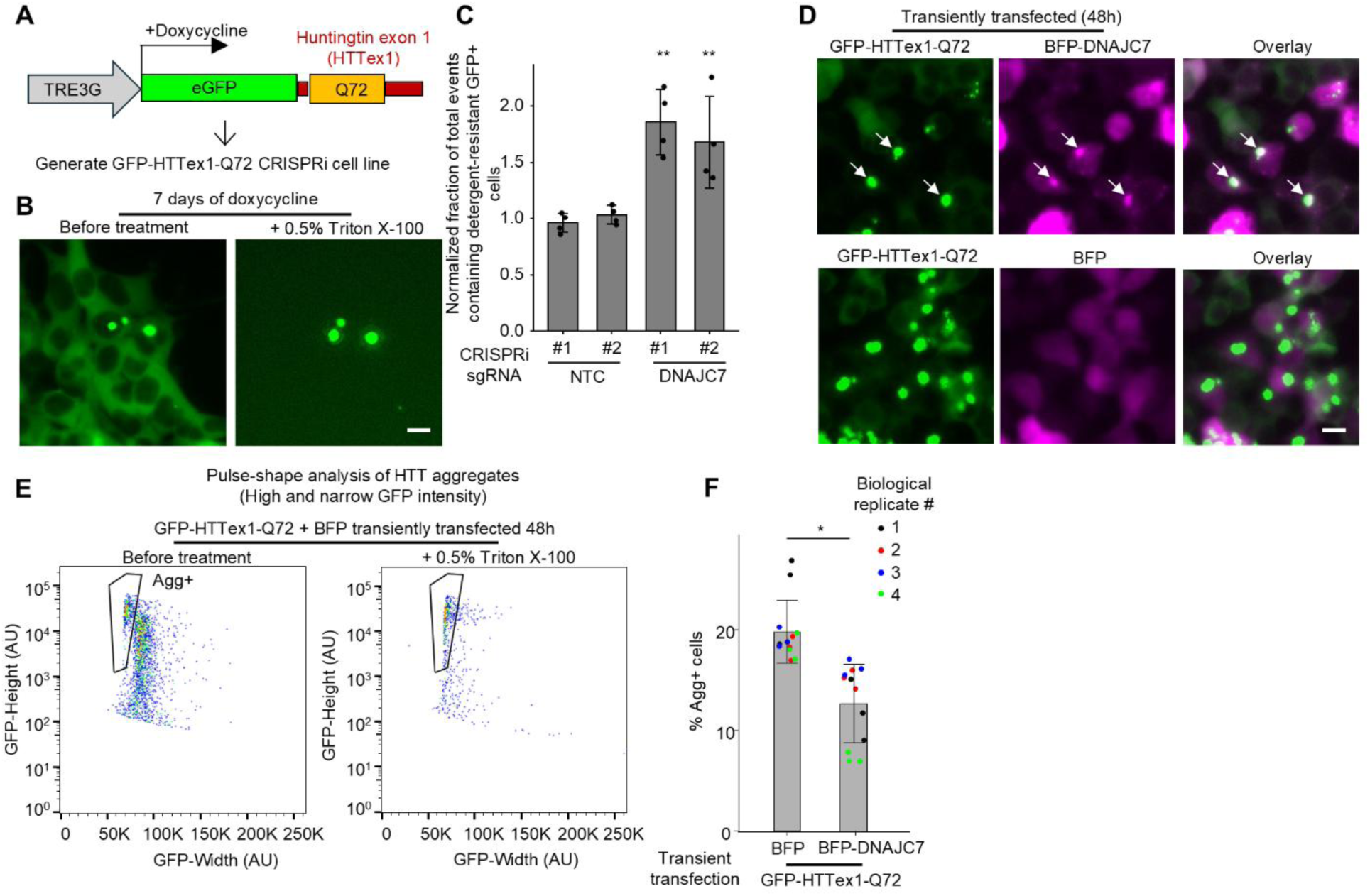
An inducible model of polyglycine (polyG) aggregation reveals detergent-resistant, p62-positive nuclear inclusions and seeding activity. **A**, Schematic of lentiviral constructs for generating the NLS-FRET-G100 cell line expressing the upstream open reading frame (uORF) of the *NOTCH2NLC* gene with a polyglycine tract of 100 residues. **B**, Immunofluorescence staining for p62 of NLS-FRET-G100 cells at 5 days of doxycycline. **C**, Fluorescence imaging of NLS-FRET-G100 cells at 5 days of doxycycline before and after detergent treatment, in the same field of view. **D**, Flow cytometry plots at one and five days of doxycycline treatment, and the latter before and after detergent treatment. **E**, Flow cytometry plots of NLS-FRET-G100 cells at 4 days of doxycycline with or without proteasomal inhibitor (carfilzomib). **F**, Flow cytometry results of NLS-FRET-G100 cells at 3 days of doxycycline with and without transfection with NLS-FRET-G100 cell homogenates (collected five days after doxycycline) for 48 h. All scale bars are 10 µm.

Together, these findings establish NLS-FRET-G100 as a robust live-cell reporter of nuclear polyG aggregation that recapitulates key pathological features of NIID and enables quantitative, high-throughput analysis.

### PolyQ and polyG proteins aggregate independently in a homotypic repeat-dependent manner

To determine whether aggregate formation by each FRET reporter is driven by the specific amino acid repeat rather than nonspecific aggregation of fluorescent tags or other aggregation-prone proteins, we co-transfected cells with different combinations of mScarlet- or mNeonGreen-tagged Q24 or Q79 constructs, uN2C-G12 or uN2C-G100 constructs, as well as GFP-HTTex1-Q25 or -Q72. The nuclear localization signal was removed from all constructs to promote predominantly cytosolic localization, as HTTex1 is known to localize primarily to the cytoplasm. We observed that HTTex1-Q72 inclusions sequestered both Q24 and Q79, but not uN2C-G12 proteins (Fig. 5). Conversely, uN2C-G100 inclusions sequestered uN2C-G12 proteins despite the short repeat length, but did not recruit Q24, HTTex1-Q25, or HTTex1-Q72 proteins. In all cases, the fluorescent proteins alone were not recruited into inclusions. The sequestration of proteins containing non-expanded polyQ repeats by polyQ aggregates has been reported previously [17,43–46]. Recent work has similarly shown sequestration of endogenous polyG-containing proteins such as FAM98B into polyG aggregates [47].

**Fig. 5.**
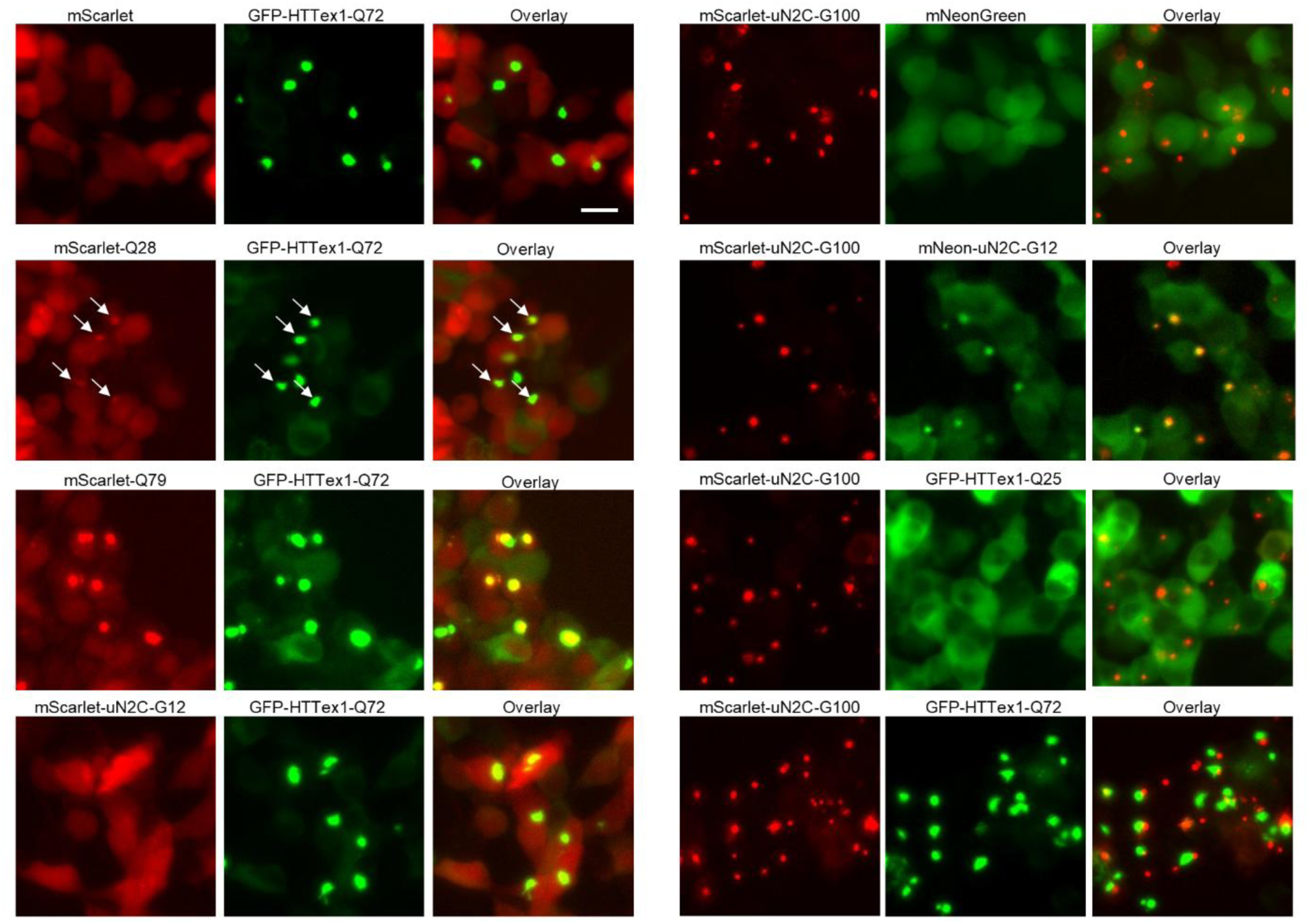
PolyQ and polyG proteins undergo independent, repeat-dependent aggregation. Fluorescence micrographs of HEK293T cells co-transfected with the indicated combinations of fluorescently tagged polyQ and polyG proteins. Scale bar: 10 µm.

Interestingly, in cells containing both HTTex1-Q72 and uN2C-G100 inclusions, aggregates were most frequently adjacent to each other, while only some showed partial overlap. Similar spatial segregation was reported previously between HTTex1 and poly(glycine-alanine)dipeptide repeat aggregates [48]. In sum, these findings confirmed aggregation in the FRET reporter is driven in a homotypic amino acid repeat-dependent manner and can lead to sequestration of other proteins containing even relatively short repeats of the same amino acid.

We also tested whether polyQ or polyG aggregates from homogenized cells or tissues could seed aggregation of the FRET reporter. NLS-FRET-Q79 cells transfected with homogenates of NLS-FRET-Q79 cells or from cortical tissue of transgenic HD mice led to an increase in FRET-high cells in (Fig. S5). We used 22-week-old mice of the R6/1 HD mouse model, which expresses exon 1 of human huntingtin (HTTex1) with approximately 115 glutamines and exhibits widespread aggregation of mutant HTT throughout the brain beginning as early as 8 weeks of age [49,50]. Immunostaining the contralateral hemispheres of these mice confirmed frequent p62-positive nuclear inclusions in the cortex (Fig. S6). NLS-FRET-G100 cell homogenates did not increase FRET-high cells in the NLS-FRET-Q79 line but robustly increased it in the NLS-FRET-G100 cells. These results indicate that the FRET reporter lines could potentially be used as a “biosensor” to test the presence of polyQ aggregates of other cells, tissues, or biofluids as for FRET-based aggregation reporters of Tau and TDP-43 [19–21,51–53].

### Chaperone screening of polyG aggregation shows few modifiers but polyG aggregation is suppressed by DNAJC7 overexpression

Using the same CRISPRi screening approach as with the NLS-FRET-Q79 cell line, we performed a molecular chaperone screen in the NLS-FRET-G100 reporter line to identify modifiers of polyG aggregation (Fig. 6A). Surprisingly, this screen revealed relatively few significant hits. Notably, key polyQ modifiers such as *DNAJC7*, *DNAJB6*, *DNAJB1*, and *HSPA8* were not identified as hits in the polyG screen. A direct comparison of Gene Scores between the polyQ and polyG screens revealed minimal overlap in chaperone modifiers (Fig. 6B), suggesting distinct mechanisms of proteostasis regulation between polyQ and polyG aggregates.

**Fig. 6.**
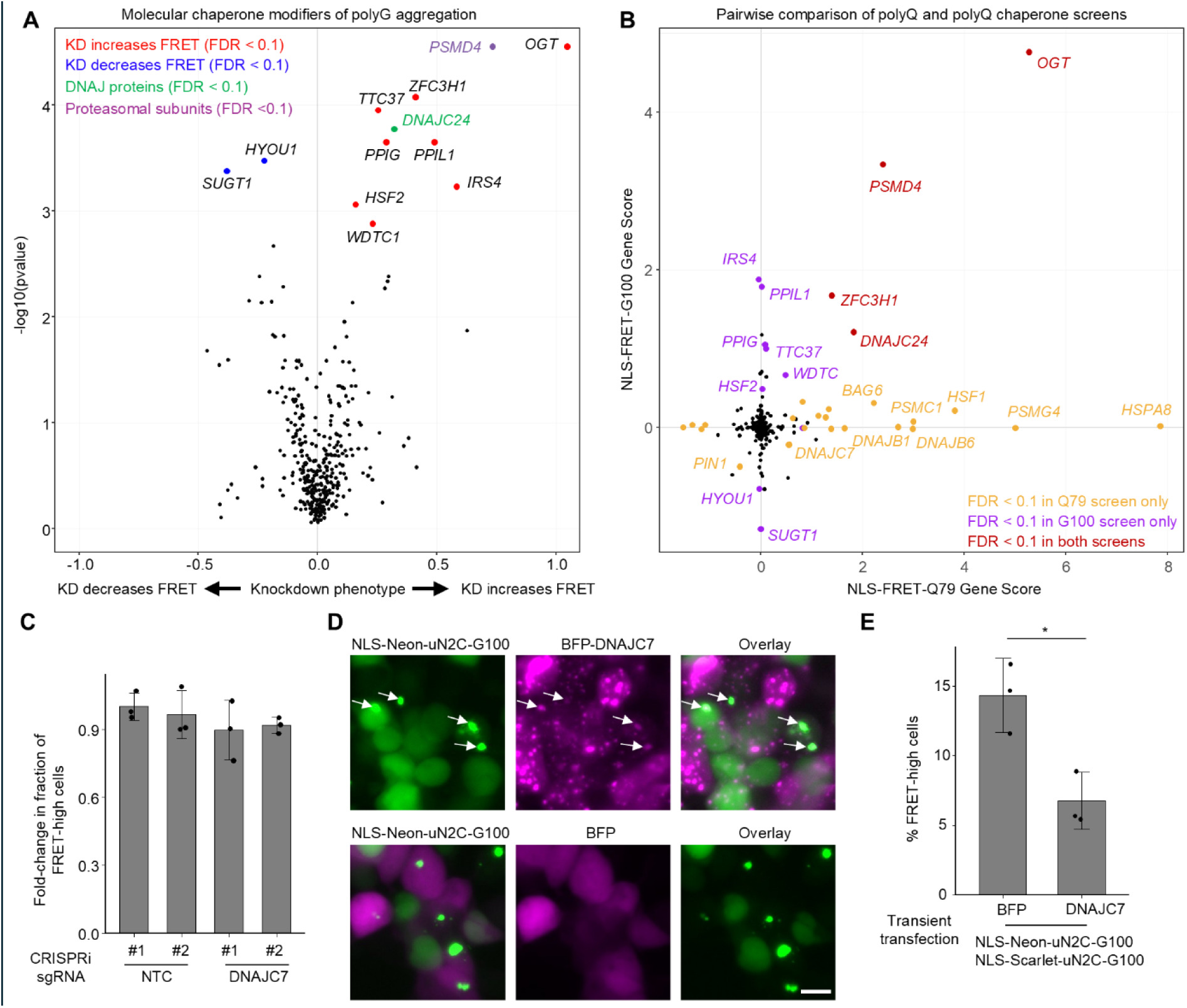
Chaperone CRISPRi screening reveals limited suppression of polyG aggregation despite effects by DNAJC7 overexpression. **A**, Volcano plot showing results of a CRISPRi screen for molecular chaperone modifiers of polyG aggregation in the NLS-FRET-G100 cell line. **B**, Pairwise comparison of Gene Scores between the polyG screen (from panel **A**) and the polyQ screen (Fig. 2B). **C**, Results of flow cytometry experiments measuring fold-change in the fraction of FRET-high cells after 5 days of doxycycline in sgRNA^+^ cells transduced with NTC or DNAJC7. Data represent mean ± sd from *n* = 3 independent experiments. **D**, Representative fluorescence micrographs of HEK293T cells transiently co-transfected with NLS-mScarlet- and NLS-mNeonGreen-tagged uN2C-G100 together with either BFP or BFP-DNAJC7; only the mNeonGreen channel is shown. Scale bar: 10 µm. **E**, Flow cytometry quantification of the percentage of FRET-high cells relative to total cells expressing both mNeonGreen and mScarlet. Bars indicate mean ± sd from *n* = 3 independent experiments. *p < 0.05 by Student’s *t*-test.

Among the few shared hits was *OGT*, which encodes O-GlcNAc transferase. *OGT* knockdown was among the top hits in both screens, and we independently validated that its depletion significantly increases the fraction of FRET-high cells in both the NLS-FRET-Q79 and NLS-FRET-G100 lines (Fig. S7), confirming the reliability of the polyG screen and confirming active CRISPRi machinery. In contrast, direct knockdown of *DNAJC7* had no significant effect on the proportion of FRET-high cells in the NLS-FRET-G100 model (Fig. 6C).

To determine whether this difference was associated with a lack of DNAJC7 colocalization with polyG inclusions, we transiently transfected cells with NLS- and mScarlet-and mNeonGreen-tagged uN2C-G100 together with either BFP or BFP-DNAJC7, as described above. Similar to observations with GFP-HTTex1-Q72 inclusions, a subset of polyG inclusions colocalized with BFP-DNAJC7 (Fig. 6D). Consistent with this finding, flow cytometry revealed a significant reduction in the fraction of FRET-high cells (Fig. 6E). We repeated these experiments using a non-nuclear mScarlet/mNeonGreen-tagged uN2C-G100 and observed the same results (Fig. S8).

To exclude the possibility that the observed effects of DNAJC7 knockdown were due to clonal artifacts of the selected FRET reporter lines, we examined one additional polyQ and multiple other polyG monoclonal cell lines. Consistent with results obtained with the NLS-FRET-Q79 and HTTex1-Q72 lines, DNAJC7 knockdown robustly increased aggregation in the polyQ line (Fig. S9). In contrast, analysis of three additional uN2C-G100 monoclonal lines with comparable FRET signal properties, two of which lacked nuclear localization, revealed no effect of DNAJC7 knockdown on aggregation in any clone. Together, these data support that polyG aggregation is insensitive to DNAJC7 knockdown, whereas DNAJC7 overexpression can reduce polyG aggregation.

## Discussion

In this study, we developed inducible FRET-based reporter systems for polyQ and polyG aggregation that recapitulate features of human disease and enable dynamic monitoring of aggregation. Importantly, these reporters allow for high-throughput flow cytometry-based genetic screens, which we used to systematically test all known molecular chaperones to identify modifiers of aggregation. Beyond screening applications, these models also provide a valuable platform for exploring the relatively understudied nuclear proteostasis pathways, and they could serve as biosensors to assess seeding activity of mouse or human brain homogenates or other biospecimens.

By taking a comprehensive approach, we were able to identify for the first time the Hsp40 co-chaperone DNAJC7 as a potent modifier of polyQ protein aggregation and further found that DNAJC7 overexpression suppresses both polyQ and polyG aggregation. A major next step is to understand the functional impact of DNAJC7 in the central nervous system (CNS), particularly given its high expression among Hsp40 family members. DNAJC7 appears to be especially important in motor neurons, supported by its genetic link to amyotrophic lateral sclerosis (ALS) and recent evidence that loss of DNAJC7 in iPSC-derived motor neurons increases susceptibility to proteotoxic stress, in part due to impaired HSF1 signaling [54].

However, its role may extend more broadly across CNS diseases. Prior studies have shown that DNAJC7 interacts with other aggregation-prone proteins, including Tau and TDP-43, with Hou et al. demonstrating that mutant tau co-precipitates with anti-DNAJC7 antibodies in a transgenic mouse model of frontotemporal dementia [31]. These findings suggest that DNAJC7 may act as a general regulator of protein aggregation in neurodegenerative disease. Our study was limited by the availability of an antibody that could reliably immunostain DNAJC7 to test its colocalization in mouse or human brain tissues to further validate this finding.

The growing number of possible DNAJC7 substrates raises questions regarding the basis of its broad substrate selectivity. As previously demonstrated for DNAJB6, DNAJC7 may preferentially bind to fibrillar or amyloid-like structures, a biophysical state shared by tau and polyQ aggregates [55–57]. To date, the amyloidogenic potential of polyG aggregates is not defined. Future studies using live-cell imaging approaches such as fluorescence recovery after photobleaching (FRAP), along with biochemical characterization could help determine if amyloidogenic state is required for DNAJC7 binding, as well as the dynamics of its recruitment. It is notable that DNAJC7 is structurally distinct among Hsp40 family members, as it contains multiple tetratricopeptide repeat (TPR) domains in addition to the canonical J-domain. Beyond the shared J-domain, DNAJC7 has no significant sequence homology with DNAJB6 or other Hsp40s, suggesting that it may recognize and engage substrates through a different mechanism. Although we observe physical interaction between DNAJC7 and polyQ proteins in cells, it remains unclear whether this reflects direct binding to misfolded species or occurs through intermediary factors. As previously shown with tau [31,58], biochemical reconstitution studies using purified proteins will be critical to determine whether DNAJC7 directly recognizes misfolded polyQ and polyG aggregates, and to dissect the specific contribution of its TPR domains to client binding and chaperone activity.

The basis for the relative insensitivity of polyG aggregation to knockdown of endogenous molecular chaperones, despite clear colocalization with DNAJC7 and reduced aggregation upon DNAJC7 overexpression, remains unclear. Discrepancies between knockdown and overexpression phenotypes are well documented [59,60]. DNAJC7 can seemingly engage polyG aggregates, consistent with recent proteomic analyses of the insoluble fraction of uN2C-polyG immunoprecipitated proteins that identified DNAJC7 as being significantly enriched [47]. One possibility is that endogenous levels of DNAJC7 may be insufficient to counteract the robust aggregation propensity of polyG proteins. Indeed, we observed that the formation of detergent-resistant, FRET-high aggregates in polyG reporter lines was markedly more robust than in comparable polyQ reporters. Testing shorter, yet aggregation-prone, polyG repeats that more closely match the aggregation properties of the polyQ reporters could reveal an effect. Alternatively, differences in aggregation kinetics between polyG and polyQ proteins may mean that the time points examined in this study were not optimal to capture the effects of chaperone depletion.

The FRET-based reporter lines described here, to our knowledge, represent the first monoclonal cell lines to exhibit spontaneous protein aggregation, even in the absence of exogenous seeding. Despite being clonal, there is a seemingly stochastic emergence of microscopically visible aggregates in only a subset of cells, the basis of which remains unclear and represents an intriguing avenue for investigation. To an extent, this phenomenon parallels observations in human disease and transgenic mouse models, where visible aggregates are present in only a subset of neurons within otherwise homogeneous populations. In our FRET reporter lines, cells lacking detectable aggregates nonetheless retain the capacity to aggregate, as aggregation can be accelerated by genetic perturbations, proteasome inhibition, or exogenous seeding. Together, these features make the reporter lines a tractable system for interrogating intrinsic cellular factors that govern spontaneous aggregate formation. In addition, given the relatively low fraction of FRET-high cells in the polyQ reporter (typically single-digit percentages), ongoing efforts include increasing expression levels, expanding repeat length, and slowing cell proliferation to enhance aggregation and improve the throughput of large-scale genetic screens for modifiers.

Beyond DNAJC7, our CRISPRi screens uncovered several additional candidate modifiers of protein aggregation that merit further investigation. One of the most prominent was OGT, the enzyme responsible for O-GlcNAcylation, whose knockdown markedly increased aggregation in both the polyQ and polyG models. O-GlcNAcylation is a dynamic post-translational modification that regulates a wide array of proteins and cellular processes, including in the brain [61–64], with our findings suggesting a potential role in proteostasis. Another notable hit specific to polyQ protein aggregates was BAG6, a chaperone “holdase” required for ubiquitin-mediated degradation of newly synthesized misfolded polypeptides that has previously been shown to modulate the aggregation of TDP-43 fragments [65–67].

Additionally, hits unique to the polyG screen included members of the peptidyl-prolyl isomerase family, such as *PPIG* and *PPIL1*, which have been implicated in modulating tau aggregation [68]. These findings highlight the potential of our platform to identify both shared and distinct regulators of aggregation across different disease-relevant proteins.

In summary, this work establishes a robust and scalable platform for studying the aggregation of polyQ and polyG proteins, leveraging inducible FRET-based reporters that enable high-throughput genetic screening in live cells. Our CRISPRi screens provide new insights into the molecular chaperone network and its role in regulating aggregation, uncovering both shared and distinct modifiers between two different repeat expansion disorders. Given the scalability of this system, ongoing efforts to extend these screens to a genome-wide level are expected to uncover additional regulators, offering deeper understanding of proteostasis mechanisms and informing the development of targeted therapeutic strategies for neurodegenerative diseases.

## Experimental procedures

### Animals

All mice were maintained according to the National Institutes of Health guidelines and all procedures used in this study were approved by the UCSF Institutional Animal Care and Use Committee. Mice were housed on a 12-h light/dark cycle at 22-25 °C, 50-60% humidity, and had food and water provided ad libitum. The mice used in this study were R6/1 mouse model of HD (B6.Cg-Tg(HDexon1)61Gpb/J, RRID:IMSR_JAX:006471), which are crossed onto a homozygous for Cre-inducible CRISPR interference machinery (B6;129S6-Gt(ROSA)26Sortm2(CAG-cas9*/ZNF10*)Gers/J, RRID: IMSR_JAX:033066). For genotyping HD mice, genomic DNA was purified from ear punches following the manufacturer’s instructions (New England Biolabs, T3010S). 1 µL of the eluted DNA was used for PCR amplification using primers and cycling conditions detailed in Supplementary Table 1.

### Plasmid construction and lentivirus packaging

A list of all plasmids used in this study with information on their main expression elements is provided in Supplementary Table 1. A map of each plasmid in a GenBank file format is provided in a Supplementary File. The FRET-based aggregation reporters were constructed on a lentiviral pLEX-TetOne backbone kindly provided by Michael Ward (National Institutes of Health) for doxycycline-inducible expression under a Tet-ON 3G promoter. C-terminal ataxin-3 (amino acid 257 to the C-terminus) containing a 24 or 79 consecutive glutamine stretch was cloned by PCR from pEGFP-Ataxin3Q28 or Q84, respectively [69] (Addgene #22122 and #22123, gifts from Henry Paulson). Huntingtin exon 1 (HTTex1) with Q25 and Q72 repeats were cloned from pGW1-HTTN586 constructs [70], kindly provided by Steven Finkbeiner (Gladstone Institutes). The upstream ORF of the *NOTCH2NLC* encoding 12 or 100 glycines [42] (Addgene #224355 and #224356, gifts from Nicolas Charlet-Berguerand) were cloned by restriction digestion into the pLEX-TetOne backbone. mNeonGreen, mScarlet, and eGFP were cloned by PCR. A 2 × c-myc nuclear localization signal and the mTagBFP2 (BFP) sequences were PCR amplified from pMK1334 [60]. The cDNA for DNAJC7 was kindly provided by Dr. Jason Gestwicki (UCSF). BFP and BFP fused to DNAJC7 were cloned by PCR into pMK1200 [71], a lentiviral backbone with an EF1α promoter.

Transient transfection experiments that used GFP-HTTex1-Q72 and 3×FLAG-tagged constructs were built on an AAV backbone with an EF1α promoter [72] (Addgene #55636, a gift from Karl Deisseroth).

The CRISPRi molecular chaperone sgRNA library used in this study was described previously [73]. Individual sgRNAs for validation studies were selected and cloned into pMK1334 lentiviral plasmid backbone between *BstXI* and *BlpI* by annealing and ligating annealed oligonucleotides. The sgRNA protospacer sequences are provided in Supplementary Table 1.

Lentiviral packaging was performed as previously described [74]. Briefly, HEK293T cells were seeded in complete DMEM to reach approximately 70% confluency the following day. For transfection, third-generation lentiviral packaging plasmids (pRSV, pMDL, and pVSV-G) were mixed at a 1:1:1 mass ratio (lentiviral pack-mix) and combined with an equal mass of the transfer plasmid. The DNA mixture was diluted in Opti-MEM and complexed with polyethylenimine (PEI; Polysciences, 23966) at a 3:1 PEI:DNA mass ratio. After 15 min of incubation at room temperature, the transfection mixture was added dropwise to the cells.

Conditioned media was collected 48 h post-transfection and filter-sterilized using a Millex-GV syringe filter unit (Millipore, SLGV033RB). Lentivirus was then precipitated using the Alstem Lentivirus Concentration Kit (VC100) according to the manufacturer’s instructions and resuspended in 1× DPBS (Sigma-Aldrich, D8537).

For pooled sgRNA library packaging, 15 µg of sgRNA library DNA and 15 µg lentiviral pack-mix) were transfected into HEK293T cells plated in a 15 cm plate format. The precipitated virus was resuspended in 5 mL of 1× DPBS. For individual sgRNAs cloned into the pMK1334 backbone, 1 µg of the transfer DNA and 1 µg lentiviral pack-mix were transfected onto HEK293T cells plated in a 6-well (35mm) plate format. The resulting virus was resuspended in 200 µL of 1× DPBS.

### Cell culture and cell line generation

All cells were maintained in a tissue culture incubator (37 °C, 5% CO2) and checked regularly for mycoplasma contamination. HEK293T cells were cultured in DMEM supplemented with 10% fetal bovine serum (Seradigm 89510-186), Penicillin-streptomycin (Gibco, 15140122), and L-glutamine (Life Technologies, 25030081).

The doxycycline-inducible NLS-FRET-Q79 and NLS-FRET-G100 cell lines were made on the “cXG284” HEK293T cell line that has stably integrated CRISPRi machinery (dCas9-BFP-KRAB) in the CLYBL locus [75]. The cells were transduced with lentivirus and, after at least 72 h and in the presence of 2 µg/ml doxycycline, the cells were dissociated and sorted for those expressing both mNeonGreen and mScarlet using a BD FACSAria FusionCell Sorter.

Monoclonal lines were obtained by plating cells at limiting dilution, followed by screening for cells that showed the highest frequency of inclusions by microscopy and by corresponding presence of a population of cells with higher FRET signal. The GFP-HTTex1-Q72 monoclonal cell line was generated similarly but used HEK293T cells expressing dCas9-BFP-KRAB introduced by random integration of lentivirus.

### Transfections, fluorescence imaging and immunofluorescence

All transfections of HEK293T cells were conducted the next day after plating into wells by diluting plasmids into Opti-MEM containing PEI at a 3:1 PEI:DNA mass ratio, incubated at room temperature for 15 min, and dispensed dropwise over the cells. The well format and amount of DNA transfected for the different experiments are detailed below.

Live-cell fluorescence imaging was performed using an ECHO Revolve microscope with a 20× objective. To assess detergent sensitivity, cells were first imaged under baseline conditions. Triton X-100 (5% stock in 1× DPBS) was then added directly to the well to achieve a final concentration of 0.5%, followed by a 1-min incubation. The same field of view was subsequently re-imaged to assess detergent-resistant fluorescence.

For immunocytochemistry of p62, cells were fixed at room temperature for 10 min with 4% paraformaldehyde (Electron Microscopy Sciences, 15710) diluted in 1× DPBS, then briefly rinsed with 1× DPBS. Cells were permeabilized and blocked for 10 min in blocking solution consisting of 1×DPBS with 0.1% Triton X-100 and 5% normal goat serum. Primary antibody against p62 (clone D5L7G, Cell Signaling, 88588) was diluted 1:1000 in blocking solution and incubated overnight at 4°C. The next day, cells were washed three times with 1× DPBS and incubated with a secondary antibody, goat anti-mouse Alexa Fluor 647, for 1h at room temperature (ThermoFisher, A32728). Nuclei were counterstained with Hoechst 33342 (ThermoFisher, 5553141) at a 1:2000 dilution. p62 antibody specificity was validated by confirming absence of signal with sgRNA-mediated knockdown.

For immunostaining of mouse brain tissue, the right hemispheres were drop-fixed in 4% paraformaldehyde overnight at 4°C, then cryoprotected in 30% sucrose prepared in 1× DPBS for at least 24 h. Brains were sectioned at 40 µm thickness, and free-floating sections were blocked in 1× DPBS containing 0.3% Triton X-100 and 5% normal goat serum for 1 h at room temperature. Sections were then incubated overnight at 4°C with a rodent-specific anti-p62 antibody (clone D6M5X, Cell Signaling, 23214) at a 1:500 dilution in blocking buffer. The next day, slices were washed three times for 10 min each in 1× DPBS and incubated with goat anti-rabbit Alexa Fluor 488 secondary antibody (ThermoFisher, A11008) for 1 h at room temperature. Nuclei were counterstained with Hoechst 33342 (1:2000 dilution in 1× DPBS) for 5 min, followed by three additional 10-min washes in fresh 1× DPBS. Sections were then mounted onto microscope slides (Fisher Scientific, 12-550-143) and coverslipped using ProLong Gold antifade mounting medium (Invitrogen, P36930).

To examine expression and co-localization of polyQ and polyG aggregates with different fluorescent tags, cells plated in 96-well format were transfected with 200 ng of each indicated polyQ, polyG, or control plasmid and with the addition of doxycycline to 2 µg/ml, followed by imaging at 48h to 72h.

Fluorescent images of all immunostained cells and tissues were acquired on the ECHO Revolve on a 20× objective.

### Primary CRISPRi screen and analysis

For screening with the chaperone pooled sgRNA library, 15 million cells containing the FRET reporter were transduced with 1ml of lentivirus and plated on a T175 flask (Day 0). On Day 2, with ∼30% of the cells showing BFP positivity, the cells were passaged and replated with 2 μg/ml puromycin (Gibco, A1113803). This was repeated on day 5. On day 7, with > 70% of cells showing BFP positivity, the cells were passaged and 20 million cells were plated without puromycin and with the addition of 2 µg/ml doxycycline. The cells were regularly passaged until day 5. The cells were dissociated with trypsin, resuspended in complete DMEM, and sorted using the BD FACSAria Fusion Cell Sorter into FRET-low (∼6 million cells) and FRET-high (∼2 million cells). Sorting and gDNA isolation for the polyQ chaperone screen was performed twice (two separate times from the same starting population of library-transduced cells). Sorting for the polyG chaperone screen was performed once. Genomic DNA was isolated using a Monarch gDNA extraction kit according to manufacturer protocols. sgRNA-encoding regions were then amplified, followed by sequencing of the protospacers by Illumina NextSeq2000 as recently described [76].

The generation of knockdown phenotypes, p-values, and gene scores to identify the hits used bioinformatics pipelines that we recently described [76], including ‘sgcount’ (https://github.com/noamteyssier/sgcount) and ‘crispr_screen’ (https://github.com/noamteyssier/crispr_screen/). In brief, raw sequencing reads were aligned to a custom reference file containing the CRISPRi chaperone library protospacer sequences using ‘sgcount’, generating sgRNA count matrices for each sample. Using ‘crispr_screen’, these counts were then normalized, and p-values were calculated for individual sgRNAs based on differential abundance between FRET-low and FRET-high. Gene-level phenotypes, including knockdown phenotypes, p-values, and false discovery rates, were derived using the Robust Rank Aggregation (RRA) algorithm [77]. The sgRNA counts matrices and gene-level phenotypes are provided in Supplementary Table 2.

### Secondary assays based on flow cytometry

NLS-FRET-Q79 or NLS-FRET-G100 cell lines were seeded (300,000 cells/well) into 6-well format and transduced with lentivirus packaged from sgRNA-containing pMK1334. After at least 72 h, reporter expression was induced by doxycycline and analyzed after 5 days by flow cytometry using a BD FACS Fortessa, specifically examining BFP^+^ (sgRNA-containing) cells. Biological replicates of the experiments were from passaging the transduced cells. Gating strategies for representative experiments are shown in Fig. S5.

For assessing detergent-resistant aggregates by flow cytometry, we first acquired 10,000 events to determine baseline fluorescence intensity. We then added 5% Triton X-100 diluted in 1× DPBS directly into the tube to a final concentration of 0.5% Triton X-100, gently swirled the tube, and performed flow cytometry to collect another 10,000 events.

Detergent-resistant GFP^+^ aggregates in the eGFP-HTTex1-Q72 cell line shown in Fig. 4 were evaluated by flow cytometry after 7 days of doxycycline treatment. The relative fold-change in fraction of FRET-high cells was obtained by normalizing to sgRNA NTC#1 condition for each experiment.

To test the effect of BFP-DNAJC7 overexpression on GFP-HTTex1-Q72 aggregates HEK293T cells (seeded the day before at 1 × 10^5^ cells/well in 24-well plate) were transfected with 500 ng of EF1⍺-driven BFP or BFP-tagged DNAJC7 and 250 ng of GFP-HTTex1-Q72 or only 250 ng of GFP-HTTex1-Q72 After a 15 min incubation, the mixture was placed dropwise over cells of one well. After 48h, we obtained fluorescence micrographs, followed by dissociating the cells for flow cytometry. We used pulse shape analysis (plotting GFP-Height versus GFP-Width) to quantify aggregates.

To test the effect of BFP-DNAJC7 overexpression on polyG FRET-high aggregates, HEK293T cells were seeded as above and transfected with plasmids (250 ng each) expressing doxycycline-inducible mScarlet and mNeonGreen-tagged uN2C-G12 or uN2C-G100 (with or without an NLS) were co-transfected with BFP or BFP-DNAJC7. 2 µg/ml doxycycline was added at the time of transfection. Imaging to examine co-localization and flow cytometry for FRET were conducted 48h after transfection.

### Transfecting cell or tissue homogenates for seeding aggregation

NLS-FRET-Q79 cells containing 2 µg/ml doxycycline were grown in a 10 cm plate for 5 days. The cells were dissociated with trypsin, spun down at 200 g x 10 min, and resuspended in 200 µL 1× DPBS. For tissue homogenates, we used fresh frozen left hemispheres of 22-week-old R6/1 male transgenic mice or age-matched non-transgenic mice. All mice are homozygous for conditional CRISPRi machinery. The right hemispheres of the same mice were drop-fixed in 4% PFA, cryoprotected, and immunostained as described above. The left hemispheres were frozen directly in dry ice and stored at -80°C. A portion of the fresh-frozen cortex was placed in 200 µL 1×DPBS and homogenized in a 1.5 ml microfuge tube. The crude homogenates from cells or tissues were briefly sonicated using a Q700 Sonicator (QSonica Sonicators) with a 1/16” microtip probe at amplitude 10 for 10 s. The samples were centrifuged at 20,000g x 15 min at 4°C, followed by collection of the supernatant into a new tube. The concentrations of the homogenates were estimated by Nanodrop absorbance at 280 nm, using 1 Abs = 1mg/ml. 100 µg of homogenates were diluted into 50 µL Opti-MEM containing 2 µL PEI (1mg/ml), and dispensed dropwise over NLS-FRET-Q79 cells plated in a 24-well format that have been treated with 2 µg/ml doxycycline for one day.

### Immunoprecipitation and western blot

To evaluate DNAJC7 knockdown, NLS-FRET-Q79 cells were transduced with lentivirus containing sgRNAs targeting DNAJC7 or non-targeting controls for 48 h. sgRNA-expressing cells were selected using 2 µg/ml puromycin for 3 days, followed by an additional two days of recovery. At 7 days post-transduction, cells were plated in a 12-well format to confluency, using 3 wells per condition. The following day, the cells were lysed with 100 µL of RIPA with protease inhibitors, sonicated briefly, and centrifuged at 21,000 × g for 15 min. The supernatant was collected and combined with LDS buffer (Invitrogen, B0007) and 50 mM DTT (Cell Signaling, 7016) and boiled for 5 min.

3 µL of each lysate was resolved on a 4-20% Tris-Glycine gel (BioRad, 4561096), transferred to a 0.2 µm nitrocellulose membrane (BioRad, 1620112) using a Trans-blot Turbo, and blocked for 1 h with Intercept® Blocking Buffer (LicorBio) before incubated with primary antibody at 4°C overnight in the same buffer. The next day, secondary antibody was incubated for 1h room temperature. Membranes were washed 3x 5 min with 1x TBS with 0.1% Tween.

Primary antibodies include rabbit anti-DNAJC7 (Proteintech, 11090-1-AP) 1:2000 and mouse anti-GAPDH (Proteintech, 60004-1-Ig) 1:2,000. Secondary antibodies are IRDye® 800CW Goat anti-Rabbit (LicorBio, 926-32211) and IRDye® 700CW Goat anti-Mouse (LicorBio, 926-68070). To assess total resolved protein, the blot was stripped with 1× NewBlot IR Stripping Buffer (LicorBio, 928-40028) for 15 minutes, washed for 5 mins with 1× TBS 3 times, stained with Revert 700 Total Protein Stain (LicorBio, 926-11011) for 10 mins. Membranes were imaged on the Licor Odyssey, and band densities were quantified using ImageJ.

### Graphing and statistical analysis

All bar graphs, scatter plots, and volcano plots were generated using R. Statistical tests were performed in R as indicated in the figure legends, except in CRISPR screening analysis which is described separately.

## Acknowledgements

The authors would like to thank members of the Kampmann Lab for discussions on data analysis and the manuscript figures. We also Lin Yadanar and Molly O’Brien for assistance with maintaining the mouse lines used in this study. We are grateful to Jason Gestwicki, Corey Nadel, Oleta Johnson, Saugat Pokhrel, and Wyatt Powell for materials and discussions related to DNAJC7. We thank the UCSF Laboratory for Cell Analysis for assistance with cell sorting.

## Author contributions

BR and MK contributed to the study’s overall conception, design, and interpretation. BR created the figures. BR and MK wrote the manuscript, with KE contributing to the writing of a component of Materials and Methods. BR and KE conducted experiments.

## Funding and additional information

This work was supported by the National Institutes of Health and National Institute of Neurological Disorders, including grants R25NS070680 (BR) and 1K08NS133300 (BR), the Larry L. Hillblom Foundation (BR), and a Ben Barres Early Career Acceleration Award from the Chan Zuckerberg Neurodegeneration Challenge Network (MK).

## Conflict of interest

BR and MK have filed a patent application on in vivo screening methods. MK is an inventor on US Patent 11,254,933 related to CRISPRi and CRISPRa screening, a co-scientific founder of Montara Therapeutics and serves on the Scientific Advisory Boards of Engine Biosciences, Alector, and Montara Therapeutics, and is an advisor to Modulo Bio and Theseus Therapeutics.

## Abbreviations and nomenclature

ALS: Amyotrophic lateral sclerosis
CRISPRi: CRISPR interference
FRET: Föster resonance energy transfer
HD: Huntington’s disease
HTTex1: Huntingtin exon 1
NIID: Neuronal intranuclear inclusion disease
NLS: Nuclear localization signal
OGT: O-GlcNAc transferase
PolyG: Polyglycine
PolyQ: Polyglutamine
TDP-43: TAR DNA-binding protein 43
TPR: Tetratricopeptide repeat
uN2C: Upstream open reading frame of *NOTCH2NLC*

## Figures and figure legends

**Fig. S1.**
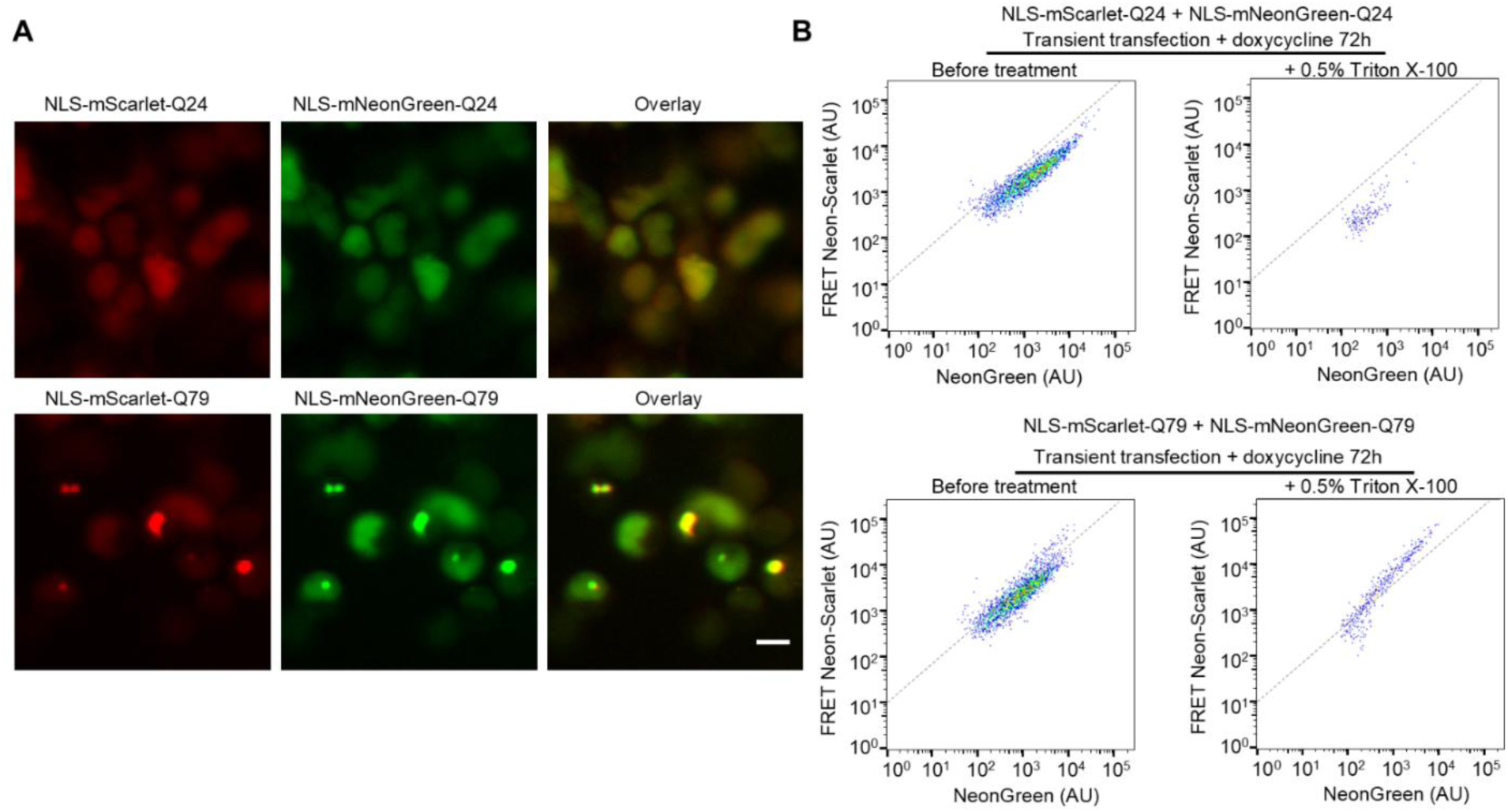
Repeat length dependence of polyQ protein aggregation and FRET signal. A, Fluorescence micrographs of HEK293T cells co-transfected with FRET-paired Q24 or Q79 constructs. B, Flow cytometry results at 72h after transfection, before and after treatment of the sample with detergent.

**Fig. S2.**
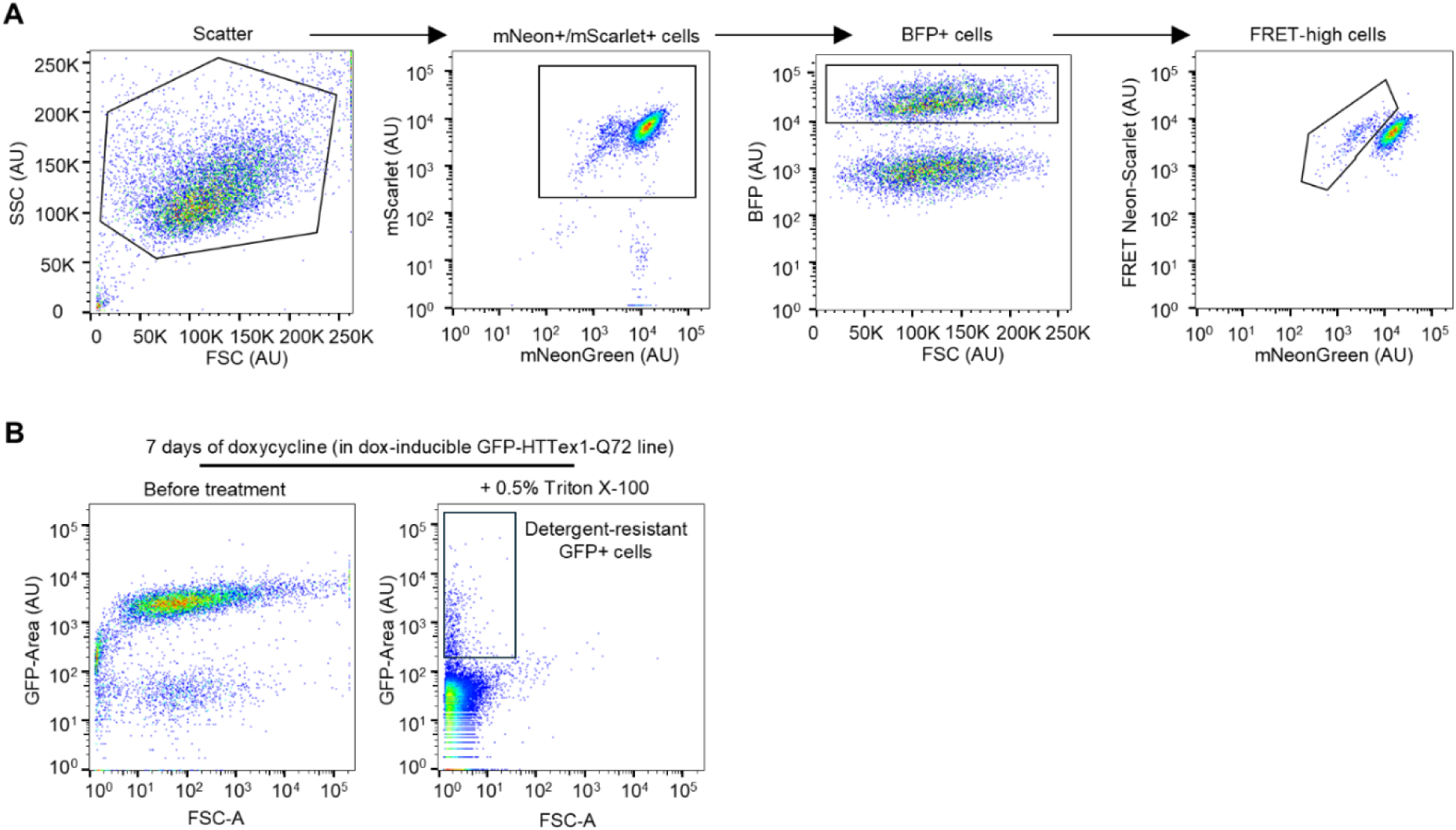
Representative gating strategies for FRET in sgRNA transduced cells and for detergent-resistant HTTex1-Q72 GFP+ aggregates. A, Example of the flow cytometry gating strategy to assess fraction of FRET-high cells of the NLS-Q79-FRET at 5 days of doxycycline treatment and transduced with a non-targeting control (NTC) sgRNA. The sgRNA expression vector contains a nuclear mTagBFP2 fluorescent reporter. B, example gating of the GFP-HTTex1-Q72 CRISPRi cell line shown in Fig. 4 transduced with NTC sgRNA.

**Fig. S3.**
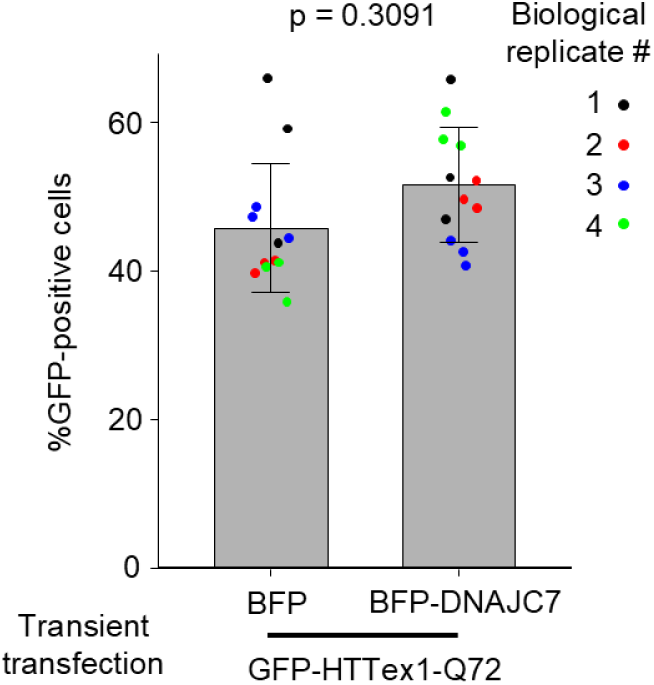
BFP-DNAJC7 does not reduce the number of GFP-positive cells when co-transfected with GFP-HTTex1-Q72. Bar graph plotting number of GFP+ cells between cells transfected with BFP or BFP-DNAJC7 corresponding to the same samples as Fig. 4f. Bars and error bars represent mean ± sd. p-value shown is by Student’s t-test.

**Fig. S4.**
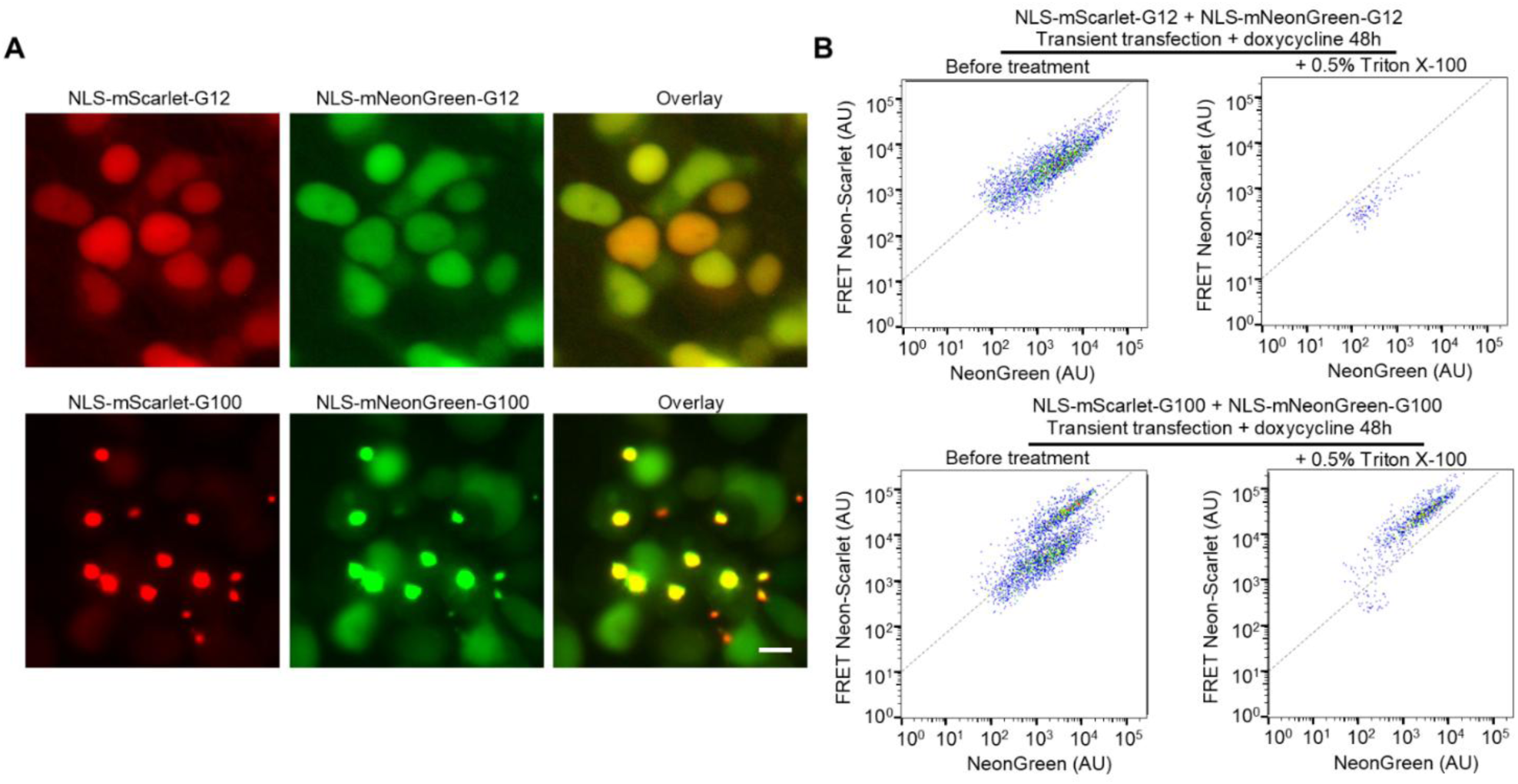
Repeat length dependence of polyG protein aggregation and FRET signal. A, Fluorescence micrographs of HEK293T cells co-transfected with fluorescent protein-tagged uN2C-G12 or –G100 constructs. B, Flow cytometry results at 72h after transfection, before and after treatment of the sample with detergent.

**Fig. S5.**
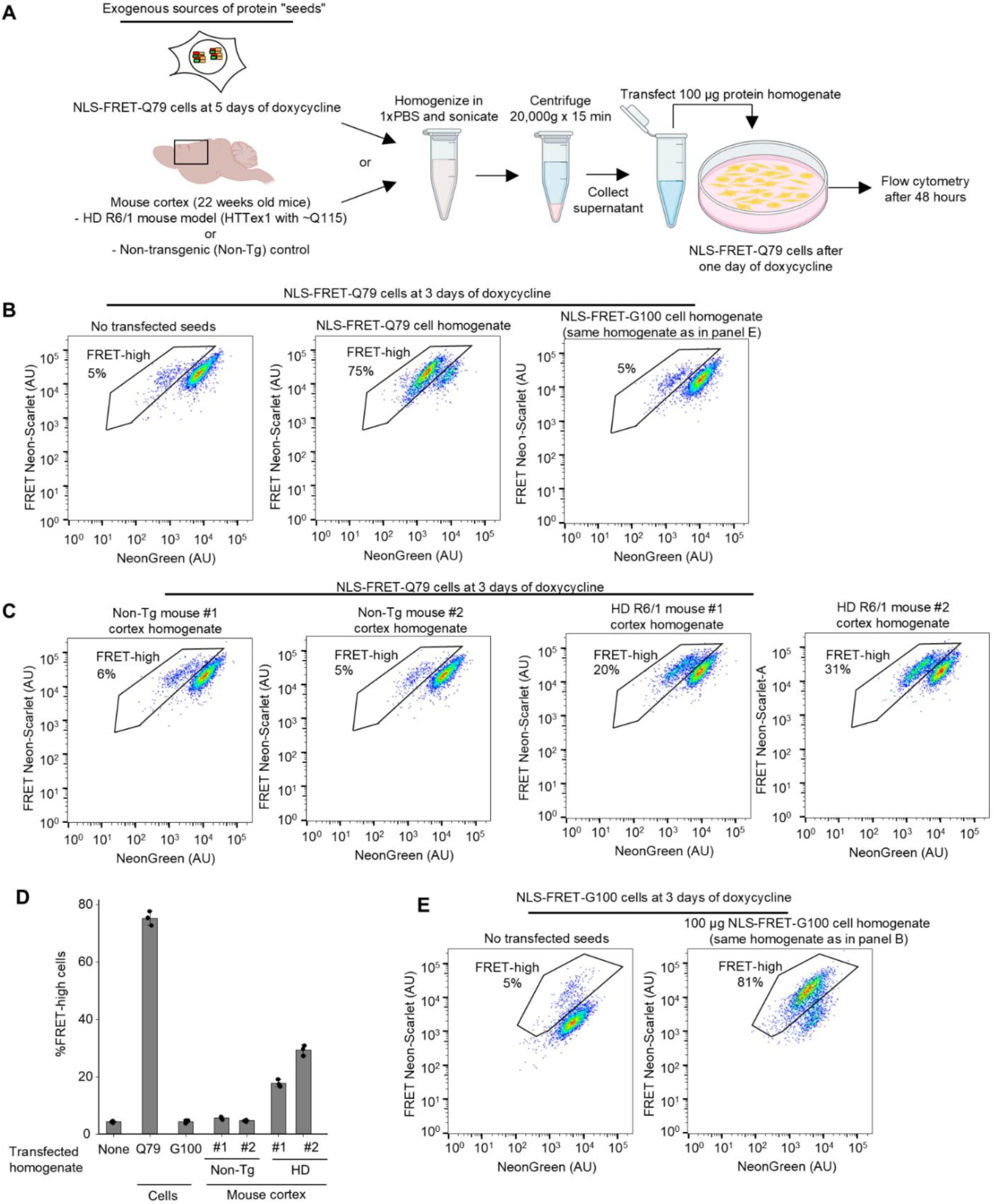
Seeding aggregation of polyQ and polyG FRET reporters with exogenous protein aggregates. A, Workflow for generating homogenates from cultured cells or mouse cortical tissue, followed by transfection into the NLS-FRET-Q79 reporter cell line. Created in Biorender. B-D, Flow cytometry plots showing fraction of FRET-high cells in NLS-FRET-Q79 after transfection with homogenates derived from NLS-FRET-Q79 cells or NLS-FRET-G100 cells (B) and cortical homogenates from Huntington’s disease (HD) transgenic mice versus non-transgenic controls (C), with quantification of technical triplicates (D); bars and error bars represent mean ± sd. E, Flow cytometry plots showing fraction of FRET-high cells in NLS-FRET-G100 cells after transfecting homogenates of NLS-FRET-G100 cells.

**Fig S6.**
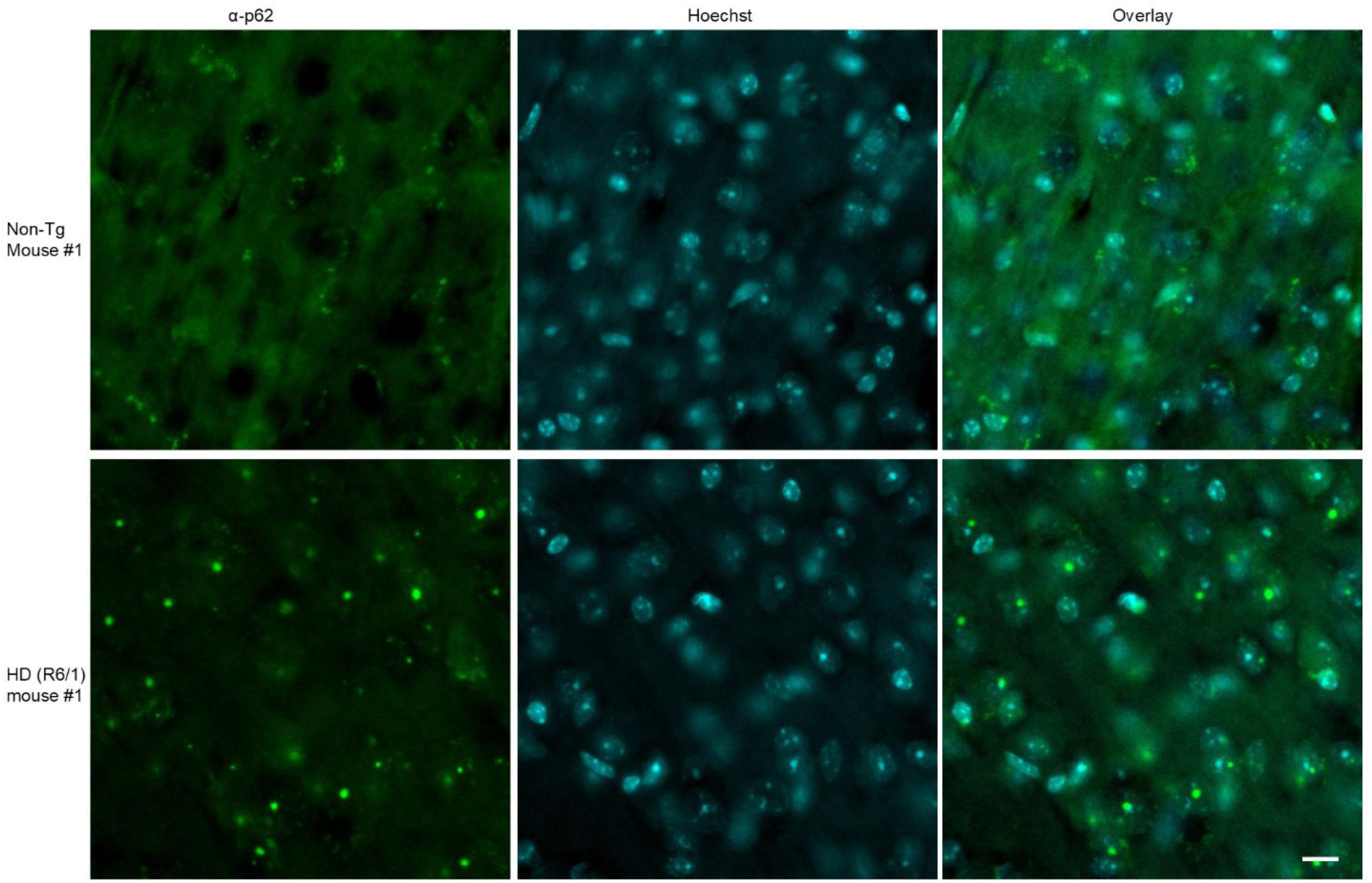
Immunostaining of HD mice shows cortical p62-positive nuclear inclusions. Representative micrographs of 40 µm brain slices immunostained for p62 (green) from a non-transgenic (top) and transgenic HD mouse (bottom), and counterstained for nuclei with Hoechst. Scale bar: 10 µm

**Fig. S7.**
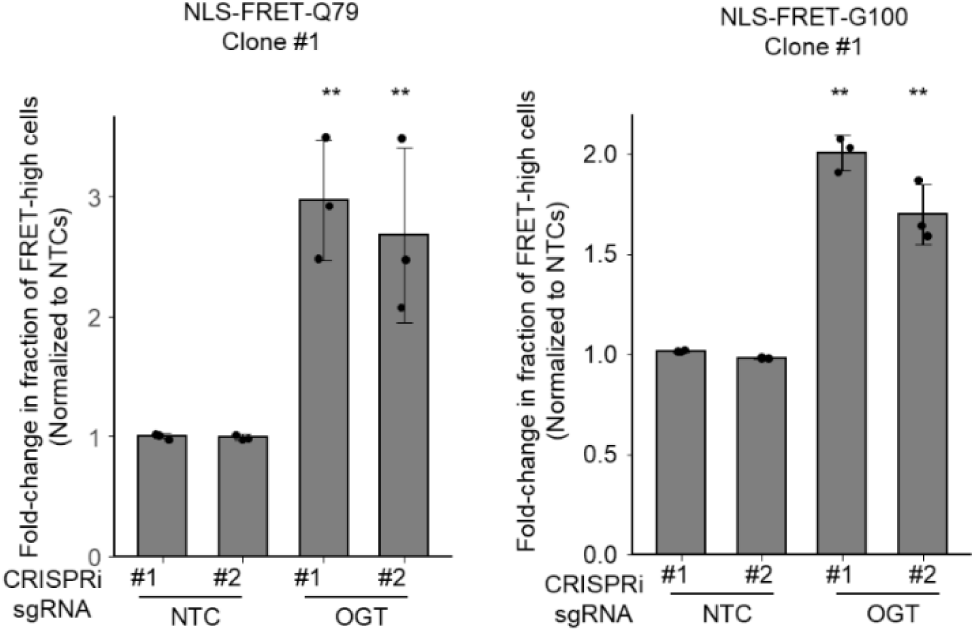
CRISPRi knockdown of OGT increases FRET in NLS-FRET-Q79 and NLS-FRET-G100 cell lines. Results of flow cytometry experiments measuring fold-change in the fraction of FRET-high cells after 5 days of doxycycline in sgRNA+ cells transduced with NTC or OGT, and in the indicated cell lines. Data show fold-change relative to the mean of the sgNTCs for each independent experiment. Bars and error bars represent mean ± sd and points represent n = 3 independent experiments. **p < 0.05, one-sample t-test compared to sgNTCs (fold-change = 1.0)

**Fig. S8.**
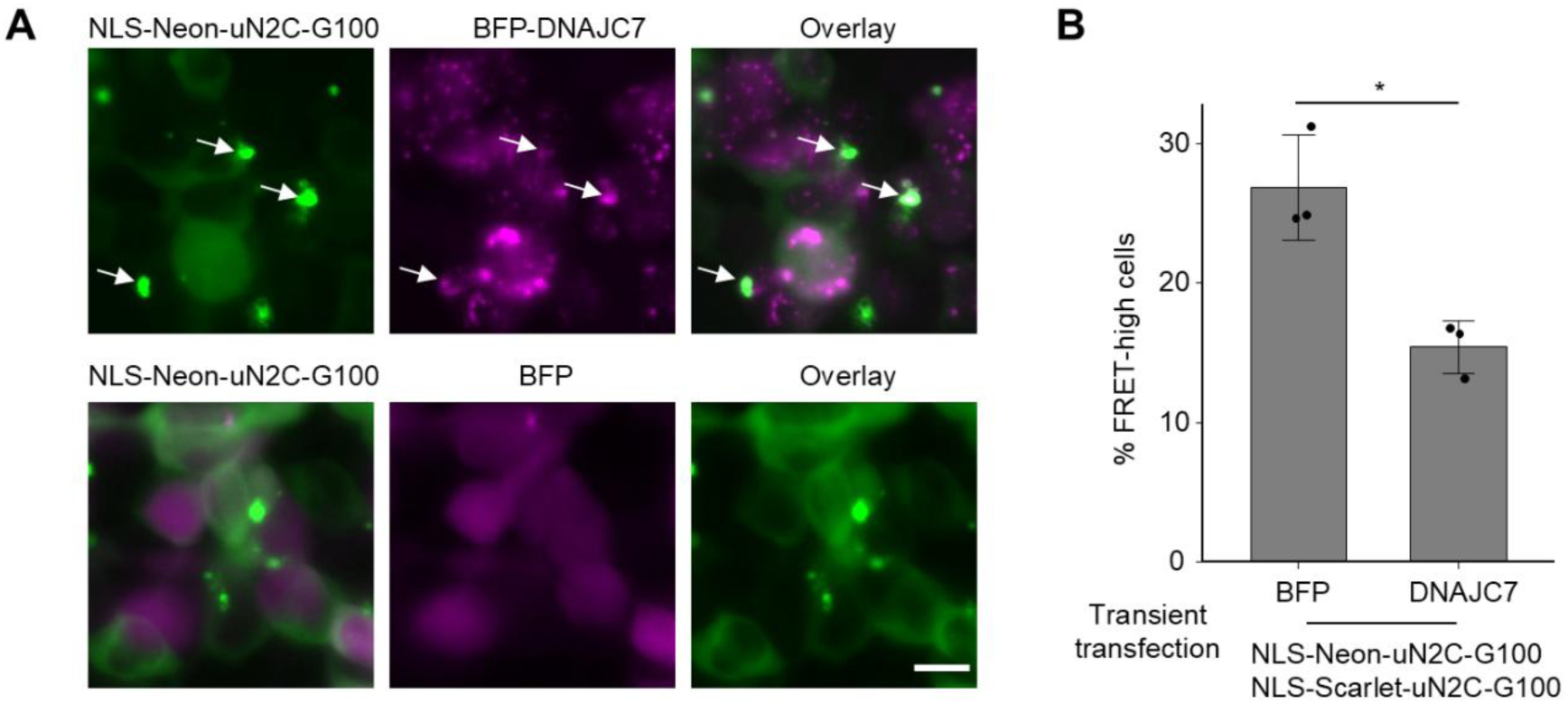
DNAJC7 overexpression reduces polyG aggregation in cells. Representative fluorescence micrographs of HEK293T cells transiently transfected with mScarlet– and mNeonGreen–tagged uN2C-G100 together with either BFP or BFP-DNAJC7; only the mNeonGreen channel is shown. Flow cytometry quantification of the percentage of FRET-high cells relative to total cells expressing both mNeonGreen and mScarlet. Bars indicate mean ± sd from n = 3 independent experiments. *p < 0.05 by Student’s t test.

**Fig. S9.**
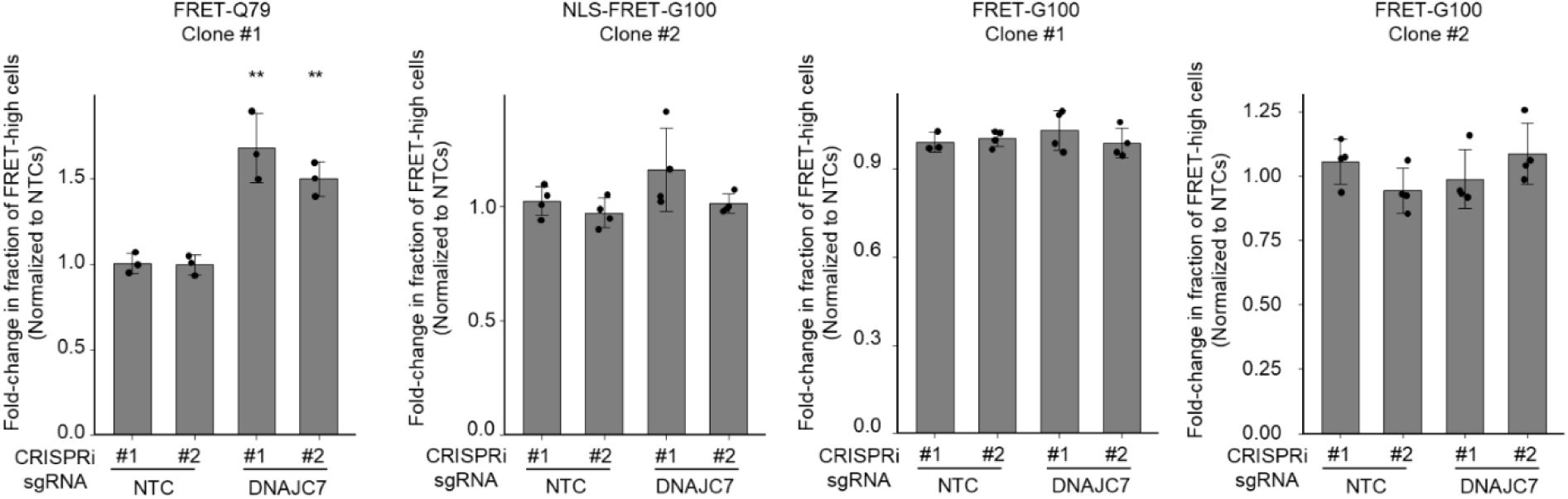
CRISPRi knockdown of DNAJC7 fails to increase FRET in multiple G100 cell lines. Results of flow cytometry experiments measuring fold-change in the fraction of FRET-high cells after 5 days of doxycycline in sgRNA+ cells transduced with sgNTC or sgDNAJC7, and in the indicated cell lines. Data show fold-change relative to the mean of the sgNTCs for each independent experiment. Bars and error bars represent mean ± sd and points represent n = 3 independent experiments. **p < 0.05, one-sample t-test compared to sgNTCs (fold-change = 1.0)

## Notes

### Summary of Updates

Control experiments showing that non-expanded polyQ and polyG proteins do not aggregate to confirm repeat length-dependent aggregation and FRET signal; added DNAJC7 overexpression data to show modifying effect of polyglyine proteins; data showing homotypic repeat-dependent aggregation added; supplemental tables updated. Co-IP data removed; updated manuscript title; updated results and discussion

