## Supplementary Table Legends for "The Hsp40 co-chaperone DNAJC7 regulates polyglutamine aggregation and exhibits context-dependent effects on polyglycine aggregation"

**Supplementary Table 1.** Genotyping primer sequences, sgRNA protospacer sequences, and a complete list with descriptions of all plasmids used in this study.

**Supplementary Table 2. ‘**crispr_screen’ pipeline outputs and ‘sgcount’ sgRNA count matrices from molecular chaperone CRISPRi screens in NLS-FRET-Q79 and NLS-FRET-G100 cell lines. Each dataset is provided in a separate tab.
